# Global distribution and diversity of haloarchaeal pL6-family plasmids

**DOI:** 10.1101/2024.08.07.607104

**Authors:** Mike Dyall-Smith, Friedhelm Pfeiffer

## Abstract

Australian isolates of *Haloquadratum walsbyi*, a square-shaped haloarchaeon, often harbor small cryptic plasmids of the pL6-family, approximately 6 kb in size. These plasmids exhibit a highly conserved gene arrangement and encode a replicase similar to those of betapleolipoviruses. To assess their global distribution and recover more examples for analysis, fifteen additional plasmids were reconstructed from the metagenomes of seven hypersaline sites across four countries: Argentina, Australia, Puerto Rico, and Spain. Including the five previously described plasmids, the average plasmid size is 6,002 bp, with an average G+C content of 52.5%. The tetramers GGCC and CTAG are either absent or significantly under-represented, except in the two plasmids with the highest %G+C. All plasmids share a similar arrangement of genes organized as outwardly facing replication and ATPase modules, but variations were observed in some core genes, such as F2, and some plasmids had acquired accessory genes. Two plasmids, pCOLO-c1 and pISLA-c6 shared 92.7% nt identity despite originating from Argentina and Spain, respectively. Numerous metagenomic CRISPR spacers matched sequences in the fifteen reconstructed plasmids, indicating frequent invasion of haloarchaea. Spacers could be assigned to haloarchaeal genera by mapping their associated direct repeats (DR), with half of these matching to *Haloquadratum*. Finally, strand-specific metatranscriptome (RNA-seq) data could be used to demonstrate the active transcription of two pL6-family plasmids, including antisense transcripts.

## 1. Introduction

*Haloquadratum walsbyi* is an extremely halophilic archaeon (class *Halobacteria*) with cells that are thin squares. It thrives in environments with salt concentrations near saturation, such as salt lakes and saltern crystallizer ponds [1–5]. Although slow growing and relatively difficult to culture, *Hqr. walsbyi* often reaches community dominance and is the main cell type (≥ 50%) seen by microscopy [5–7] or detected by sequence analysis [8–11]. Its high community abundance has been correlated with elevated magnesium concentrations [1,2,12].

A curious feature of the genome of *Hqr. walsbyi* strain C23^T^ was the discovery of two small, circular plasmids (pL6A and pL6B) that were of similar size (6,056 and 6,129 bp) and gene content and shared 76% sequence identity [4]. Additional Australian isolates of *Hqr. walsbyi* were also found to carry pL6-related plasmids [13], indicating they were common in this genus. In an earlier metagenomic study of Lake Tyrrell, a hypersaline lake, Tully *et al*. calculated that plasmids similar to pL6B were present in at least 32-40% of the *Haloquadratum* population [9]. These plasmids were so common, that the metagenomic data from Lake Tyrrell contained sufficient reads to be able to reconstruct a pL6-like plasmid, pLTMV-6 [13]. Only five pL6-family plasmids have so far been described and all share seven core genes with a conserved synteny, a size range of 5.8-7.0 kb, and a GC content (51-53%) significantly higher than that of the chromosome of *Haloquadratum* (48%). A diagram of pL6B (Figure 1) illustrates the seven core genes arranged in two outwardly oriented modules. The replication module with forward genes F1 to F3, and the ATPase module with four reverse oriented genes R4 to R7. Only the R4 protein (Figure 1) has been experimentally detected in the proteome of *Hqr. walsbyi* [4], where it was associated with a membrane fraction, consistent with the presence of a c-terminal transmembrane domain (TMD; asterisked in Figure 1).

**Figure 1.**
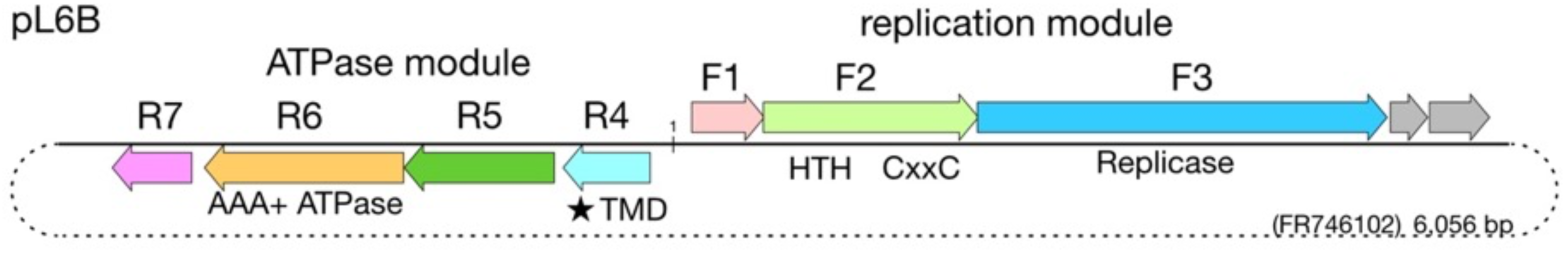
Plasmid pL6B gene diagram, showing the three core forward genes of the replication module (F1 to F3), and the four core genes of the leftwards ATPase module (R4 to R7). The plasmid start base (nt 1), shown in the center, is set between R4 and F1. HTH, helix-turn-helix domain; CxxC, Cys-x-x-Cys motif; (star)TMD, transmembrane domain. Dot-ted line indicates circularity. Sequence accession and plasmid size are given at lower right.

At the nucleotide level, these replicons are 61-79% identical. They lack integrase genes and have only been found as episomal, circular plasmids. None carry a DNA polymerase gene, but the F3 gene is proposed to encode a replicase [14,15]. In the best studied case of *Hqr. walsbyi* strain C23^T^, these plasmids are multi-copy, with the copy number of pL6A and pL6B (combined) estimated as ∼30 copies/genome equivalent [4]. It remains unclear how pL6A and pL6B are both maintained in the host cell population.

Two of the core genes of these plasmids (F3 and R6) specify proteins similar to those of haloviruses. The F3 protein of pL6B is 45% identical to the replicase protein ORF9 (HRPV-3_gp09) of Halorubrum pleomorphic virus 3 (HRPV-3), a betapleolipovirus [14,15]. R6 is a predicted AAA+ ATPase, and the pL6B protein shows 24% identity to the ATPase (His1V_gp16) of salterprovirus His1 [16,17], a lemon-shaped virus currently classified in the family *Halspiviridae* [18]. His1V_gp16 is most likely the virus packaging ATPase [19].

It is often difficult to distinguish plasmids from temperate viruses if the capsid proteins are novel, such as the case with pHK2, originally thought to be a plasmid [20] but later found to be a temperate pleolipovirus [21]. The circumstantial evidence that pL6-family plasmids may represent a novel group of temperate viruses can be summarized in three points. They carry two genes encoding proteins related to known haloviruses [13]. The seven core genes display a highly conserved synteny, and sequence conservation. They can be found in metaviromic sequence data [22], with the caveat that these post-0.1 *µ*m filtrate metagenomes probably also include DNA from extracellular vesicles (EV) [23–25]. The previously described pL6 elements vary in size from 5 to 7 kb, a range that overlaps at the top end with the smallest pleolipovirus genome (HRPV1, 7 kb), while at the low end is close to the 5,278 bp genome of Aeropyrum pernix bacilliform virus 1 (APBV1), a rod-shaped archaeal virus [26]. More generally, bacterial viruses of the *Microviridae* have DNA genomes of 5 kb and less [27].

In this study, publicly available metagenomic data from hypersaline lakes and saltern crystallizer ponds were used to reconstruct plasmids related to the known members of the pL6 family. Their annotation and comparison revealed a greater diversity in gene content and arrangement than previously recognized. These plasmids were found to match numerous CRISPR spacers retrieved from the same metagenome as well as those present in metagenomes of other hypersaline sites. In addition, recently available RNA-seq data from the microbial communities of two sites (Lake Tyrrell and Santa Pola) contained numerous reads mapping to both strands of autochthonous pL6-family plasmids, demonstrating active transcription as well as significant levels of counter-transcripts.

## 2. Materials and Methods

### Metagenomic sequence data and plasmid assembly using metaplasmidSpades

Metagenomes with *Haloquadra-tum walsbyi* were identified in the SRA database using the Branchwater server (https://branchwa-ter.jgi.doe.gov/; accessed 25/1/2024) and the *Hqr. walsbyi* C23 genome sequence (FR746099.1) as the query sequence. This returned 77 metagenomes with cANI scores of 1. These were only subjected to more detailed analysis when the read length was (a) at least 150 nt and reads were paired, or (b) average read length was >500 nt. Candidate metagenomes were individually examined for reads matching pL6-family plasmids using the SRA BLASTn option available within each SRA web page, and those showing numerous matches ( > 100) were selected for plasmid reconstruction attempts. Metagenomes from which pL6-like plasmids could be reconstructed are described in Table S1. Fastq reads were downloaded from the ENA website (https://www.ebi.ac.uk/ena/browser/home), imported into Geneious Prime v. 2023.2.1 (https://www.geneious.com) and trimmed using BBDuk (v. 38.84). Trimmed reads were exported as fastq reads and uploaded to a Galaxy server (https://usegalaxy.org.au) as input for metaplasmidSpades (with default settings).

Circular contigs assembled by metaplasmidSpades [28] were then imported back into Geneious for further analysis.

### Manual assembly of pL6-related plasmids from metagenomes

This is based on the read-extension method described earlier for reconstructing plasmid pLTMV-6 [13]. Trimmed sequence reads were first mapped to all known pL6-family plasmid sequences using parameters which relax the mapping stringency (Geneious mapper with the following settings: custom sensitivity, fine tuning, none(fast/read mapping), max mis-matches/read = 35%, max ambiguity =5%, allow gaps (2%), max gap size = 6). Reads matching the F3 genes were extracted, de-novo assembled using stringent assembly parameters (Geneious assembler, custom sensi-tivity, max mismatches per read = 2%, max ambiguity = 2), and the resulting contigs examined for sequences of high confidence and read coverage (≥ 10x). A region (400-1200 nt) was then selected and extended at both ends by repeated rounds of read mapping using the full set of metagenomic (paired) reads at the same strin-gent assembly parameters as used for the de novo assembly. After each mapping round, the contig termini were examined for paired reads that overlapped the contig ends by at least 100 nt. In this way, contig exten-sion proceeded at high confidence. Consecutive cycles of mapping and contig extension were performed un-til the sequence repeated at the termini, indicating the borders of the plasmid had been crossed and that ring closure had been achieved. Depending on the coverage, read length and distance between read pairs, this took up to 37 rounds. A monomeric sequence was generated by trimming the terminal duplication. Subse-quently, the first base was set to a point as similar as possible to that of pL6A (accession FR746101) and the genes annotated using a combination of similarity to previously annotated pL6 plasmids, assisted by annota-tion tools including GeneMarkS-2 [29] and phanotate [30].

Plasmid pTYRR-r1 was reconstructed in two stages. In the first stage, an initial seed contig was gener-ated from RNA-seq reads (SRR24125903-4) mapped at relaxed stringency (same settings as for the other plas-mids) to the F3 gene of pLTMV-6. The mapped reads were de novo assembled with stringent assembly pa-rameters (as used for the other plasmids) and a 400 nt region of this contig was selected, based on its high confidence and read coverage. In the second stage, metaviromic DNA reads from Lake Tyrrell viral concen-trates (SRR5637210, SRR5637211, SRR402042 and SRR402044) were used to extend the RNA-seq based seed sequence by repeated rounds of read mapping (conditions same as above) until ring closure.

*CRISPR spacer searches*.Sequences from existing and newly generated plasmids were used to search for matching spacers at the IMG/VR Viral/Spacer BLAST server (https://img.jgi.doe.gov/cgi-bin/vr/main.cgi). In addition, spacers were retrieved from metagenomes using the MinCED tool (https://github.com/ctSkennerton/minced). Spacers were then imported into Geneious and formatted as a FastA database. Plasmid se-quences were used to query the spacer database and the best hits were inspected to check whether their flanking direct repeats (DR) were similar to known CRISPR arrays using the CRISPR-Cas++ database (https://crisprcas.i2bc.paris-saclay.fr), and by BLASTn searches at NCBI (https://blast.ncbi.nlm.nih.gov/) against the WGS database (Expect threshold, 10^-3^; Organism: Archaea).

### RNA-seq data used for mapping to pL6 plasmids pTYRR-r1 and pPOLA-c1

As with the metagenomic data, RNA-seq data (see Table S1) were downloaded from the ENA (https://www.ebi.ac.uk/ena/browser/home), imported into Geneious Prime (v. 2023.2.1) and the ends trimmed of low-quality bases using BBDuk (v. 38.84), available within the Geneious environment. Ribosomal RNA reads were depleted by mapping to a ribosomal RNA operon from *Hqr. walsbyi* C23 (NC_017459; nt 66,889-72,015) using the Geneious ‘map to ref-erence’ tool. Trimmed unmapped reads were mapped to plasmid sequences derived from the same hyper-saline site (pTYRR-r1/Lake Tyrrell; pPOLA-c1/Santa Pola) using the ‘map to reference’ tool within Geneious Prime (settings: custom sensitivity, fine tuning = none (fast/read mapping); max mismatches per read=1%, max ambiguity=1%; no gaps).

### Phylogenetic tree reconstructions

Complete nucleotide sequences of all pL6-family plasmids were aligned using the MAFFT (v7.490) aligner [31] within Geneious Prime (ver. 2024.0.5), and this was used to infer a consensus tree using the Geneious Tree Builder, the Neighbor-Joining method, and 1,000 bootstrap repeti-tions.

Proteins similar to the F3 or R6 proteins of pL6-family plasmids were retrieved from the Genbank nr protein database using BLASTp (blast.ncbi.nlm.nih.gov/Blast.cgi). For F3 homologs, only those that could be aligned with ≥90% of the query sequence (highest E value, 2 x 10^-54^) were selected for tree reconstructions. For R6-related proteins, only relatively few could be retrieved, and homologs with a query coverage of ≥70% were selected. The F3 proteins of the newly reconstructed pL6 plasmids were added to the proteins retrieved from GenBank, aligned using the COBALT aligner (www.ncbi.nlm.nih.gov/tools/cobalt/cobalt.cgi) with de-fault parameters, and then imported into Geneious Prime where a Neighbor-Joining consensus tree (100 bootstrap repetitions) was generated using the Geneious Tree Builder tool. The R6-related proteins were aligned using MAFFT within Geneious Prime, and then a consensus tree inferred using the Neighbor-Join-ing method (within Geneious Prime) and 1,000 boostrap replications.

### Other bioinformatics tools

Nucleotide and protein sequence alignments were performed within the Gene-ious Prime (v. 2023.2.1) environment, using the MUSCLE (v. 5.1) or MAFFT (v7.490) aligners with default settings. Protein domain searches were performed with InterProScan (https://www.ebi.ac.uk/interpro/re-sult/InterProScan) and the conserved domains server (https://www.ncbi.nlm.nih.gov/Struc-ture/cdd/wrpsb.cgi). Helix-turn-helix (HTH) motifs were detected using (https://users.cis.fiu.edu/∼giri/bio-inf/GYM2/prog.html) and (https://npsa.lyon.inserm.fr/cgi-bin/npsa_auto-mat.pl?page=/NPSA/npsa_hth.html). Tetramer analysis of DNA used (http://gscompare.ehu.eus/tools/ol-igo_frequencies/). Alphafold 2 structure predictions used Galaxy Version 2.3.1 of this software available on the galaxy server at https://usegalaxy.org.au. Protein structure similarity searches used Foldseek (https://search.foldseek.com).

## 3. Results

### 3.1. Assembly of pL6-family plasmids from publicly available metagenomes

The metagenomes of hypersaline waters used to reconstruct pL6-like plasmids are listed in Table S1, and their sample locations are indicated on the world map shown in Figure 2. These metagenomes were selected for their abundance of *Hqr. walsbyi* and high numbers of reads matching pL6-family plasmids (see Methods). The only exceptions were metavirome sequences, which contain relatively low levels of cellular DNA. Reconstruction of pL6-family plasmids from metagenome (or metavirome) reads followed two strategies. The first used multiple cycles of read extension from an initial seed sequence, and was described previously for the reconstruction of plasmid pLTMV-6 (Dyall-Smith & Pfeiffer, 2018). The second method used metaplas-midSpades, an automated pipeline [28] that assembles plasmids from metagenomic reads.

**Figure 2.**
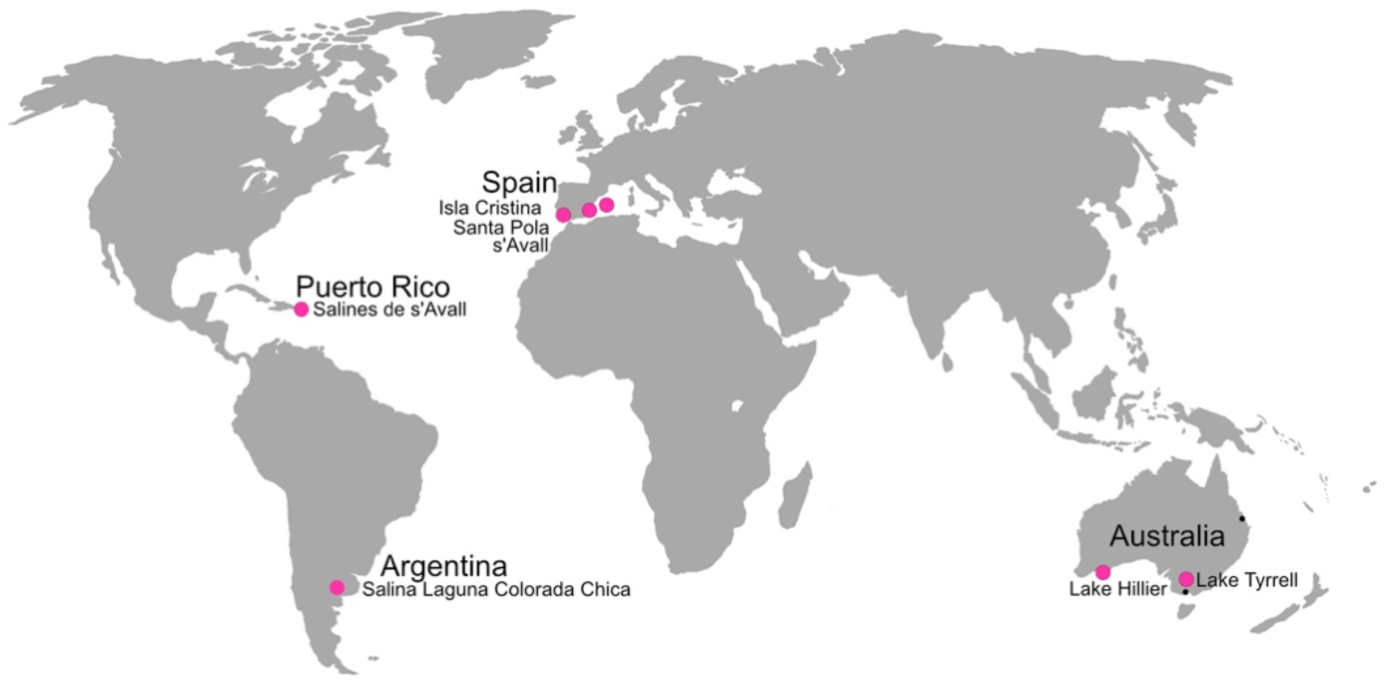
Global map of the hypersaline sites from which metagenomic data was used to reconstruct pL6-family plasmids (large pink spots). In addition, two coastal Australian sites where *Hqr. walsbyi* strains carrying pL6-family plasmids have been previously isolated are indicated by black dots.

Fifteen novel pL6-family plasmids were reconstructed and their general features are given in Table 1 along with the five previously reported pL6-family plasmids [13]. Details regarding read coverage are provided in Table S2 and Figure S1 (panels A-O). For convenience, all twenty plasmids will be used in the fol-lowing comparisons.

**Table 1.** The fifteen pL6-family plasmids reconstructed in this study along with the five previously reported plasmids (grey shaded)^a^.

| Country | Plasmid Name<sup>b</sup> | Size (bp) | %G+C | Source | Metagenome BioProject<sup>c</sup> | Plasmid Accession |
| --- | --- | --- | --- | --- | --- | --- |
| Argentina | pCOLO-c1 | 5,857 | 53.9 | Laguna Colorada Chica | PRJEB45291 | BK067852 |
| Australia | pL6A | 6,129 | 51.0 | *Hqr. walsbyi* C23 | NA<sup>d</sup> | FR746101 |
| Australia | pL6B | 6,056 | 52.0 | *Hqr. walsbyi* C23 | NA | FR746102 |
| Australia | pBAJ9-6 | 6,213 | 53.0 | *Hqr. walsbyi* Bajool9 | NA | LT984491 |
| Australia | pLT53-7 | 7,045 | 51.0 | *Hqr. walsbyi* LT53-19 | NA | LT984489 |
| Australia | pLTMV-6 | 5,884 | 53.0 | Lake Tyrrell | PRJNA81851 | LT991975 |
| Australia | pHILL-c1 | 6,187 | 52.4 | Lake Hillier | PRJNA865792 | BK067853 |
| Australia | pHILL-c2 | 7,625 | 51.5 | Lake Hillier | PRJNA865792 | BK067854 |
| Australia | pTYRR-r1 | 5,844 | 51.0 | Lake Tyrrell | PRJNA81851 | BK067859 |
| Puerto Rico | pCABO-c1 | 5,007 | 48.4 | Cabo Rojo Saltern | PRJNA343765 | BK067804 |
| Puerto Rico | pCABO-c2 | 5,889 | 53.7 | Cabo Rojo Saltern | PRJNA343765 | BK067846 |
| Puerto Rico | pCABO-c6 | 5,864 | 53.0 | Cabo Rojo Saltern | PRJNA343765 | BK067847 |
| Puerto Rico | pCABO-c9 | 6,930 | 49.3 | Cabo Rojo Saltern | PRJNA343765 | BK067848 |
| Puerto Rico | pCABO-c10 | 5,219 | 51.4 | Cabo Rojo Saltern | PRJNA343765 | BK067849 |
| Puerto Rico | pCABO-s1 | 6,206 | 47.8 | Cabo Rojo Saltern | PRJNA343765 | BK067850 |
| Puerto Rico | **pCABO-s5** | 5,017 | **58.5** | Cabo Rojo Saltern | PRJNA343765 | BK067851 |
| Spain<br>(Isla Cristina) | pISLA-c6 | 5,881 | 53.6 | Isla Cristina Saltern | PRJNA890281 | BK067855 |
| Spain<br>(Isla Cristina) | **pISLA-s1** | 5,606 | **64.0** | Isla Cristina Saltern | PRJNA890281 | BK067856 |
| Spain (Mallorca) | pMALL-c2 | 5,504 | 52.6 | s'Avall Saltern | PRJEB45291 | BK067857 |
| Spain (Santa Pola) | pPOLA-c1 | 6,075 | 48.1 | Santa Pola Saltern | PRJNA713327 | BK067858 |

<sup>a</sup>Grey shaded rows are the five plasmids reported previously [13], mostly from strains of *Haloquadratum walsbyi*. These are included to compare with the plasmids reconstructed in the present study. Bold type (pCABO-s5, pISLA-s1, and their %G+C values) indicate the two 'high GC' plasmids.
<sup>b</sup>Except for previously reported plasmids (grey shaded), plasmid names indicate source and assembly method. The suffix c is used for de novo assembled contig sequences that are extended by repeated rounds of read mapping. The suffix s denotes assembly using metaplasidSpades (see Methods). The r suffix indicates this plasmid was extended by read mapping from a contig derived from RNA-seq data.
<sup>c</sup>NCBI Bioproject accession of the metagenome read data used to reconstruct each plasmid.
<sup>d</sup>NA, not applicable, as these were sequenced from isolates not reconstructed from metagenomes.

Plasmid sizes range from 5.0 to 7.6 kb (average of 6,002 bp), and their %G+C values vary from 47.8 to 64.0 (average 52.5%, median 52.2%) but two of the new plasmids are outliers with significantly higher values than the other 18 which have %G+C values below 54%. These are pCABO-s5 (58.5%) and pISLA-s1 (64.0%). These outliers show a number of divergent features (see later) and will be referred to as high GC plasmids. DNA sequence alignment of all 20 plasmids gave an average pairwise nucleotide identity of 47.7% (range 28.5 −92.7%). The two most closely related plasmids are pCOLO-c1 and pISLA-c6 which show 92.7% nt identity, a surprisingly high value given the geographical distance between their sites of origin (Argentina and Spain, respectively).

Tetramer frequency analysis of plasmid sequences (Table S3) identified two motifs, GGCC and CTAG, with significant under-representation. GGCC is absent in 17 of the 20 plasmids and strongly under-represented in pBAJ9-6, which has only two closely spaced motifs located between the F3 and R7 genes, a common region for foreign gene acquisition (see later). However, GGCC was not under-represented in the two high GC plasmids (pCABO-s5 and pISLA-s1), indicating that this site is not under selection in their normal host species.

CTAG is absent in 10 plasmids, under-represented in 7, and present at expected levels in the two high GC plasmids (see Table S2). The remaining plasmid, pHILL-c2, was unusual in having the expected level of CTAG motifs (8 sites), but four of these sites are found between the F3 and R7 genes, all within a 122 bp region (nt 3,182-3,303) near an acquired gene encoding a formyltransferase-family protein. It is likely that the region containing these sites was also acquired, and outside this region the motif is also under-represented (0.57, second order Markov chain estimate). Finally, the tetramer TTAA was found to be absent in pISLA-s1 but present in all other plasmids.

Selection against longer motifs is more difficult to assess given the small size of pL6-family members, but the 6-mer GGATCC (e.g BamHI) is absent in all plasmids except pTYRR-r1, where it occurs only once within an acquired gene encoding a helicase that is located between the F3 and R7 genes.

The results suggest the two high GC plasmids originate from haloarchaea that do not select against CTAG and GGCC motifs, and possibly a species with a higher %G+C than *Hqr. walsbyi*. Most genera within the class *Halobacteria* have genomes with high %G+C values, ranging from 61 to 70% (see table S4 of [32]), but *Hqr. walsbyi* is atypical in having a much lower %G+C, averaging 47.8% [4].

Figure 3 shows an unrooted phylogenetic tree inferred from the complete sequences of the pL6-family plasmids (see Methods). The bootstrap confidence values at most branch points are above 95% (from 1,000 repetitions). The eight plasmids from Australian sites all branched together forming a single clade. The two high GC plasmids (left) from Spain and Puerto Rico formed a distinct clade but with long branch lengths between them. pCABO-s1 (Puerto Rico) and pPOLA-c1 (Spain) also branch closely together. The other eight plasmids show a mixture of relationships between plasmids from different sites, for example plasmids pCOLO-c1 (Argentina) and pISLA-c6 (mainland Spain) branched very closely together, and they form a sister group with pMALL-c2, from the mediterranean island of Mallorca.

**Figure 3.**
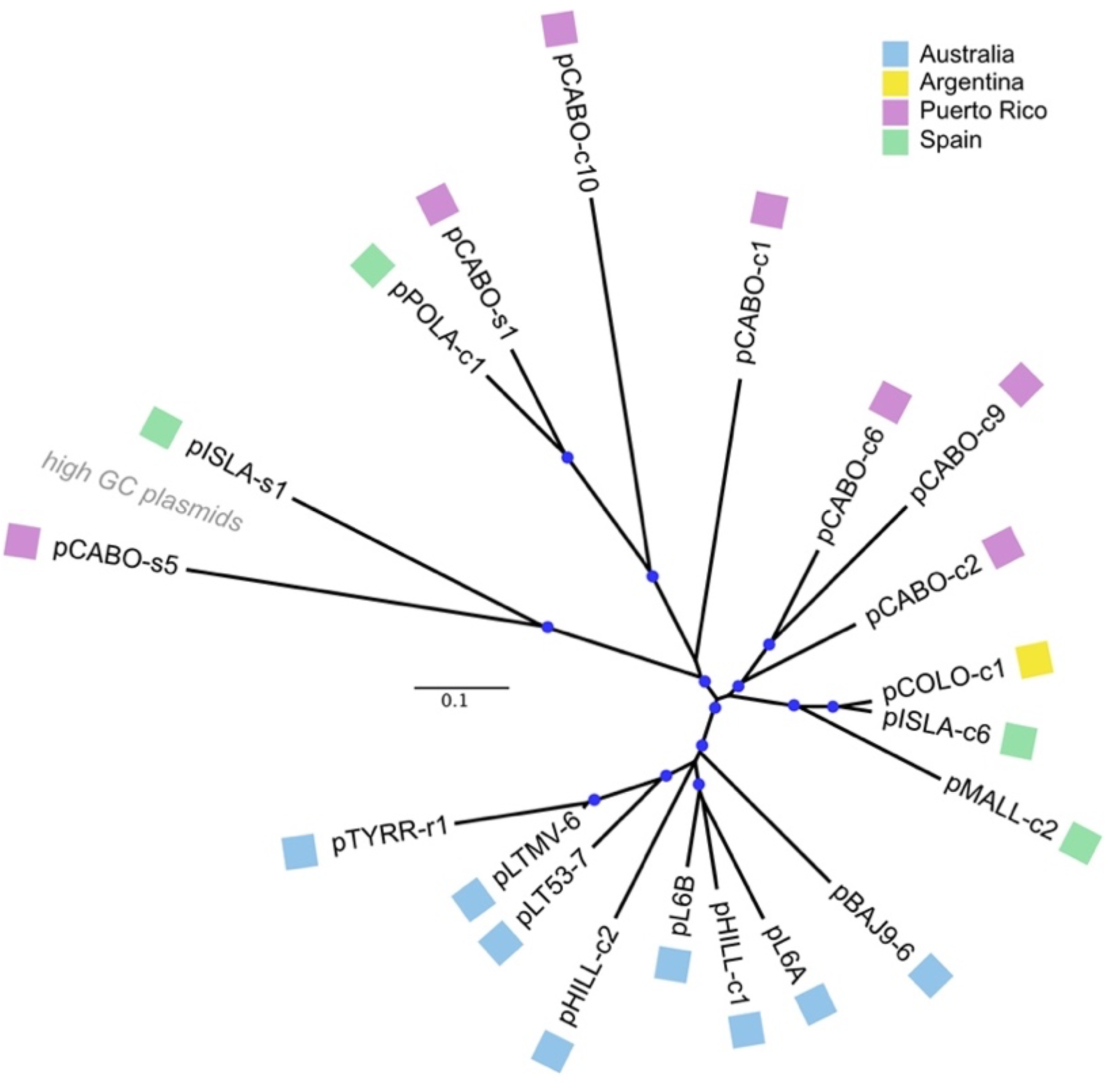
Phylogenetic tree reconstruction of pL6-family plasmids using complete nucleotide sequences (see Methods). Sequences were aligned using MAFFT and a Neighbor-Joining consensus tree generated using the Geneious Tree builder. Scale bar represents inferred changes per site. Blue circles at branch points indicate bootstrap confidence values of 96-100% (1,000 bootstrap repetitions). Colored boxes next to plasmid names show the country of origin (key at top right).

### 3.2. Relative presence of plasmid pPOLA-c1 in fractionated samples from Santa Pola crystallizer

In the study of [11], saltern crystallizer water samples from Santa Pola, Spain, were fractionated using various combinations of filtration and centrifugation to purify dissolved DNA and viruses. These brines were rich in *Haloquadratum*, with an abundance estimated at 63% of prokaryotic genera. Since pPOLA-c1 was reconstructed from the reads of that study (Table 1), it could be used to probe the relative levels of this plasmid in each fraction. Equal numbers of paired reads (3 million) from the four fractions were mapped to the pPOLA-c1 sequence and the number of matching reads for each are given in Table 2. Compared to the DNA of the cell pellet (iDNA), pPOLA-c1 was present at raised levels (1.6 −2.6-fold) in dissolved DNA produced by two methods (FdDNA, CdDNA). However, there were few matching reads in the ‘virus DNA’ preparation obtained from pelleted material collected after ultracentrifugation of cell-free fractions, suggesting pPOLA-c1 is only present as dissolved DNA. Various interpretations of these results are found in the Discussion.

**Table 2.** Relative levels of pPOLA-c1 in filtered and centrifuged fractions of Santa Pola water.^a^.

| DNA fraction | Fraction name<sup>b</sup> | Fraction description<sup>b</sup> (accession) | #Mapped reads to pPOLA-c1 | Relative proportion of mapped reads |
| --- | --- | --- | --- | --- |
| Cell pellet | iDNA | intracellular DNA (SRR13926770) | 1228 | 1 |
| Supernatant after ultracentrifugation | FdDNA | protocol F, dissolved DNA (SRR13926769) | 3217 | 2.6 |
| Supernatant after ultracentrifugation | CdDNA | protocol C, dissolved DNA (SRR13926768) | 1982 | 1.6 |
| Pellet from ultracentrifugation | vDNA | viral DNA (SRR13926767) | 39 | 0.03 |

<sup>a</sup>Metagenomic reads from the study of [11] were mapped to the sequence of plasmid pPOLA-c1 (BK067858) using the 'map to reference' tool within Geneious Prime v. 2024.0.5 (Geneious mapper, $\leq 1\%$ base mismatches). Three million paired reads were used from each of the four fractions (accessions given in the Fraction Description column). <sup>b</sup>Fraction names and Fraction Descriptions are from [11].

### 3.3. Comparative gene maps of pL6-family plasmids

The reconstructed plasmids were annotated (see Methods) and their gene maps compared to the five previously described pL6-family members (Figure 4). Plasmids are shown linearized and starting at base 1 at the left. Grey shading between corresponding genes of each map indicates their nucleotide similarity, and related CDS have been color-coded to highlight relationships. For ease of comparison, the twenty plasmids have been clustered into four groups according to the similarity of their replication modules. It is clear from Figure 4 that pL6-family plasmids share a common pattern of gene organization, although several of the newly reconstructed representatives show variations to this basic pattern which will be described below. Alignments of plasmid encoded proteins are also provided as supplementary figures (Figures S2, panels A - I) to aid these descriptions.

**Figure 4.**
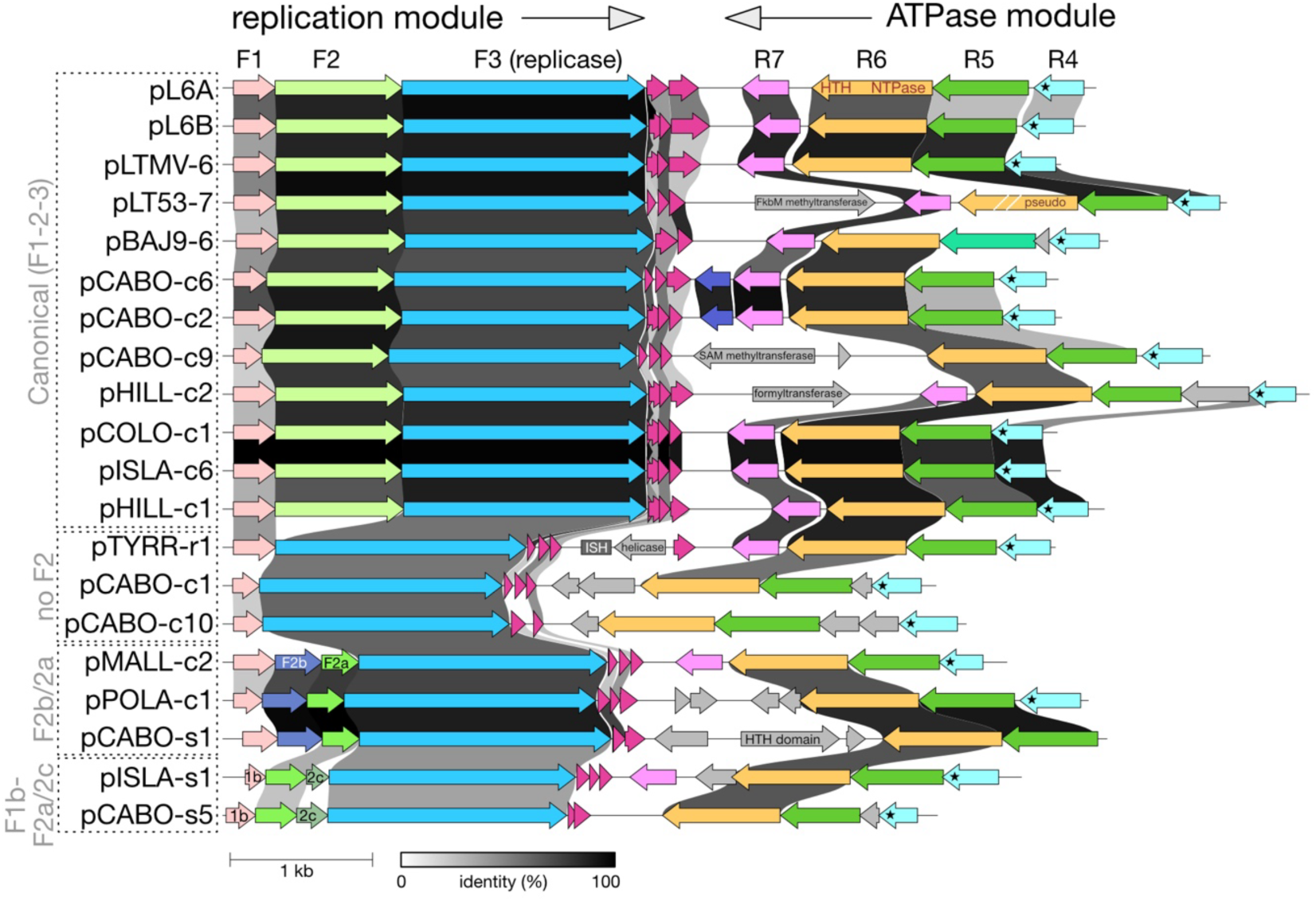
Gene maps of all pL6-family members. Plasmids are named at the left edge and have been linearized such that the starting base is leftmost. Boxed plasmid names and grey labels at the left indicate grouping of plasmids based on similarities in their replication modules (see text). Annotated genes are either color coded to indicate corresponding homologs or are grey shaded to indicate they do not have homologs in other pL6 plasmids. The previously used nomenclature of the main proteins (F1-3 and R4-7) is shown at the top. R4-proteins all have a predicted transmembrane domain (indicated by asterisks) at their C-termini. Greyscale shading between CDS indicates nucleotide similarity (see identity (%) key underneath). Scale bar (1 kb) is shown below.

#### 3.3.1 Replication (forward) module

In all plasmids, the genes from F1 to F3 in this module overlap at stop/start codons, indicating a single transcriptional unit that is probably driven by a promoter within the R4-F1 intergenic region (Fig. 1, and see later). Most plasmids (12/20) fall into the ‘canonical group’ and carry full-length versions of the three canonical forward genes F1-F2-F3 as found in pL6A. This group includes plasmids from Australia, Argentina, Puerto Rico and Spain. Deviating from this pattern are three groups of plasmids that are boxed and labeled at the left edge of the figure. The first of these, labeled ‘no F2’, consists of three plasmids that have lost the F2 gene entirely (pTYRR-r1, pCABO-c1 and pCABO-c10), indicating that this gene is either dispensable or the plasmids are defective. The next group (F2b/2a) of three plasmids (pMALL-c2, pPOLA-c1 and pCABO-s1) have replaced F2 with two smaller CDS, labeled F2b and F2a. F2a shows sequence similarity to the N-terminal region of F2 and also a structural relationship (see below). F2b is unrelated to F2. The third group (F1b/F2a/F2c) consists of the high GC plasmids pISLA-s1 and pCABO-s5, where each carries a divergent F1 gene (F1b) followed by F2a and a novel gene (F2c) before F3. F2c is unrelated to F2. In all plasmids, a number of small CDS occur just after F3 and in the same orientation (colored crimson in Fig. 4) but they will not be discussed further because they are very short and their annotation remains uncertain.

An alignment of F1 proteins (Figure S2A) shows considerable sequence diversity ranging from (a) identical or near-identical sequences (pCOLO-c1, pISLA-c6, pMALL-c2), (b) sequences showing deletions (e.g. pCABO-c6), and (c) the highly divergent F1b sequences of pISLA-s1, pCABO-s5. None contain conserved protein domains or match proteins with known function. Alphafold2 did not predict any tertiary structure. Those proteins with deletions were seen to have suffered losses within the c-terminal half, after approximately residue 50. Examination of full-length F1 gene sequences revealed the presence of direct repeats, 16-27 nt in length, that occur within the distal half, from nt 199 onwards. For example, pCOLO-c1 carries three 17 nt direct repeats of TTGCACCAACTCGTGCA between nt 277-353. These nucleotide repeats are a likely cause for deletion events within these genes.

F2 proteins have few matches in the NCBI databases (BLASTp, nr database, accessed April 22, 2024) and do not carry conserved domains that could indicate function. The twenty plasmids encode proteins that have been divided into four groups. Groups F2 and F2a represent the majority (17) and are related by sequence and structure. The F2 proteins from the twelve plasmids with canonical gene arrangement have similar lengths (296-301 aa), sequence (71-100% aa identity), and all possess a CxxC motif near the C-terminus (Figure S2B). Their alphafold2 predicted structures show two compact domains that are connected by a flexible linker (e.g. Figure S3). The five F2a proteins are much shorter (87 aa), share 31-100% aa identity to each other, and align to the N-terminal end of the canonical F2 proteins (32-50% aa identity). Their sequences encompass the first F2 domain, and alphafold2 predictions show F2a proteins fold with high confidence to form structures very similar to that of the domain 1 of F2 (Figure S3). The F2a genes of three plasmids are positioned immediately upstream of F3 while in two plasmids this gene is separated from F3 by a short, unrelated CDS (F2c, see below).

In the third group, three plasmids have an F2b gene positioned between F1 and F2a. An alignment is given in Figure S2C. The three F2b proteins are of similar length (105-109 aa) and closely related (77-97% aa identity) but show no sequence similarity to other pL6 proteins, including F2, no detectable conserved domains and no matches to proteins in GenBank (BLASTp, nr database, accessed April 22, 2024). Structural predictions using alphafold2 showed they could form a compact domain consisting of three short alpha helices and two beta strands (Figure S4) but much of this structure was of low confidence, it did not match domain 2 of F2, and structure similarity searches (https://search.foldseek.com/) did not return any significant matches.

The last group are the two F2c proteins found only in the two high GC plasmids. They are relatively short (52 and 72 aa) and share 20% aa identity (Figure S2D). The F2c of pISLA-s1 has two CxxC motifs. Neither protein retrieved matches from NCBI using BLASTp (nr database, accessed April 22, 2024).

F3 proteins are predicted to represent a novel replicase [14,15], and their alignment is shown in Figure S2E. Except for the proteins from the two high-GC plasmids (pCABO-s5, pISLA-s1) the other 18 share high sequence similarity (62-99% aa identity). The 99% aa identity of F3 proteins from pISLA-c6 and pCOLO-c1 reflects the almost identical nucleotide sequence of the replication modules of these two plasmids (99.6% nt similarity; Figure 4).

When the twenty F3 protein sequences are used to search against virus proteins (BLASTp, NCBI nr, viruses (taxid:10239), accessed 22 April 2024), the top two matches in all cases are the putative replicases of the betapleolipoviruses Halorubrum pleomorphic virus 3 (HRPV3, HRPV-3_gp09, YP_005454281) and Haloge-ometricum pleomorphic virus 1 (HGPV1, HGPV-1_gp14, YP_005454308), with 42-50% aa identity.

Overall, these comparisons of the replication module revealed that (a) F1 genes code for proteins that are diverse in sequence and can suffer deletions due to direct repeat sequences, (b) the F2 gene may be absent or be replaced by two smaller genes, one of which (F2a) represents the first domain F2, and (c) the F3 replicase is strongly conserved.

#### 3.3.2. ATPase (reverse) module

The canonical gene arrangement of R4-R5-R6-R7 is seen in the majority of plasmids (14 of 20) (Figure 4), with four of these having an additional CDS positioned either between R5 and R4 (pBAJ9-6, pHILL-c2) or between R6 and R7 (pL6A, pISLA-s1), and in the same orientation. All (20/20) carried R5 and R6 (ATPase) genes. The six plasmids with non-canonical modules lacked homologs of one or two core genes but the genes remaining always retained the same canonical order and orientation. Unlike the overlapping gene arrangement of the replication module, the core genes in this module were often not overlapping. Genes R4 and R5 either had intergenic distances of 17-38 nt (8 cases), overlapped (7 cases) or had an intervening CDS (3 cases). Genes R6 and R7 had intergenic distances of between 43-162 nt (14/20) or had inserted genes between them (3 cases). However, if atypical start codons such as CTG would be utilized then many have potential CDS that span or nearly fill these regions. The R5/R6 genes either overlapped (5 cases) or the gene distances were small (0-3 nt distance; 14/20). No plasmid showed an inserted gene between R5 and R6.

##### R4 genes

All but one plasmid carried a R4-family gene. The encoded proteins range in length from 90-134 aa and vary widely in sequence (5 −100% aa identity; Figure S2F) but all carry transmembrane domains at their C-termini, indicated by black asterisks in Figure 4. BLASTp searches of NCBI return only pL6-family homologs (accessed 22 April 2024). The single plasmid without an R4 gene (pCABO-s1) also has an unusual ATPase module that includes a foreign gene (encoding a HTH domain protein) located between genes R6 and R7 but in the opposite orientation (see below).

##### R5 genes

The predicted R5 proteins varied widely in sequence (12 −98% aa identity) but all possessed a serine as the second amino acid and all except one had the C-terminal sequence (F/Y)DYSDLI (Figure S2G). BLASTp searches of NCBI (accessed 22 April 2024) returned very few matches, with almost all being previously published pL6-family homologs. The most divergent R5 protein is that of pCABO-c10, which also lacks the C-terminal motif. The alphafold2 predicted structures of R5 proteins, such as pL6A R5 (https://www.alphafold.ebi.ac.uk/entry/G0LNF8), show a compact central domain with three alpha-helices while the C-terminal conserved motif lies at the end of a long flexible tail. The extreme conservation of the C-terminal motif across widely divergent R5 proteins indicates an important function.

##### R6 (ATPase protein) genes

All have predicted P-loop NTPase domains and C-terminal HTH domains and they share 60-99% aa identity except for the R6 protein of pCABO-c10 (Figure S2H). The pCABO-c10 protein is only distantly related to the others (19-25% aa identity) but alphafold2 predictions show it has a closely similar structure (Figure S6), with a compact P-loop ATPase domain and a helix-turn-helix domain, connected by a flexible linker. A number of similar proteins can be retrieved from the protein databases using BLASTp searches (NCBI, nr proteins) and their diversity is described further below.

R6 proteins show up to 24% aa identity with spindle-shaped halovirus His1 protein His1V_gp16 (AAQ13731), an ATPase [4,13]. An alphaFold2 structural prediction was performed to test whether their 3D structures were also similar. As shown in Figure S5 the structure of His1V_gp16 is closely similar to the R6 proteins of pL6B and pCABO-c10 (Figure S6), with the same overall architecture. Two recently discovered haloviruses also carry R6 homologs. Halorubrum spindle-shaped virus-BLv25 (OQ850971, HRSSV) encodes a protein (WLW38172, named ORF1) sharing 26% aa identity with R6 of pL6B, and Halorubrum virus V_ICIS4 (OR762182.1) specifies a protein (WPH59218, AFNJKBDN_CDS0001) that is 39% identical to pL6B R6.

##### R7 genes

Most plasmids (14/20) carried R7 genes, and their encoded proteins are closely related (61-97% aa identity; Figure S2I). BLASTp searches (NCBI, nr, accessed 22 April 2024) retrieve only the previously reported pL6-family homologs. Predicted R7 protein structures (e.g. pL6A, AF-G0LNF6-F1) show a compact arrangement of central beta-strands surrounded by three alpha-helices. Among the other six plasmids that lack an R7 gene, two carry genes in a similar position and orientation. The first is pCABO-s1 (nt 3,403-3,032) which encodes a protein with a terminal phenylalanine (as do all R7 proteins), but its overall protein similarity to R7 proteins is insignificant. The second is pCABO-c1 (nt 2,890-2,489), which instead encodes an unrelated protein matching *Haloquadratum* proteins such as Hqrw_2237.

##### Accessory genes

Some plasmids carry one or two genes that are unrelated to the core genes and appear to have been acquired by insertion into the plasmid backbone (Figure 4, grey coloring). Most are found in the region between the ends of genes F3 and R7, downstream in both cases, presumably because insertions there have little impact on the functions of replication and ATPase gene modules. Eight accessory genes (seven encoding proteins and one insertion sequence) are listed in Table S4, along with their top database matches. In all but one case, the top match was from haloarchaea, and the most common genus was *Haloquadratum*. Three proteins have general functional assignments: two methyltransferases and a formyltransferase but their natural substrates are not known.

Structural similarity of the formyltransferase enzyme specified by pHILL-c2 supported a role in sugar modification. The alphafold2 predicted structure shares close resemblance to sugar N-formyltransferases, such as dTDP-4-amino-4,6-dideoxyglucose formyltransferase from *Mycobacterium tuberculosis* (AF-P9WKZ3-F1-model_v4; foldseek E-value, 9.0 x 10^-22^; https://search.foldseek.com, accessed July 20, 2024). This enzyme converts dTDP-4-amino-4,6-dideoxyglucose into dTDP-4-formamido-4,6-dideoxyglucose and its structure has been experimentally determined [33,34]. A structural alignment with the pHILL-c2 protein is provided in Figure S7.

The gene contexts of the corresponding best matching proteins of the three transferases were examined (Figure S8, A-C). All are nearby to genes encoding enzymes involved in sugar transfer, synthesis or modification (e.g. glycosyltransferases, sialic acid synthase, polysaccharide biosynthesis protein SpsG).

### 3.4. Relationship of pL6-family F3 and R6 proteins to other homologs

Only the F3 (replicase) and R6 (ATPase) proteins retrieve significant numbers of closely related proteins from the sequence databases; 200 proteins for F3 but just 20 closely related proteins for R6 (BLASTp, NCBI, nr, accessed April 26, 2024, see Methods). All matching proteins were from haloarchaea or their viruses. The phylogenetic trees inferred from these (Figures 5 and 6) show that, with only a few exceptions, the pL6-family proteins cluster together, and away from the proteins of known haloviruses.

**Figure 5.**
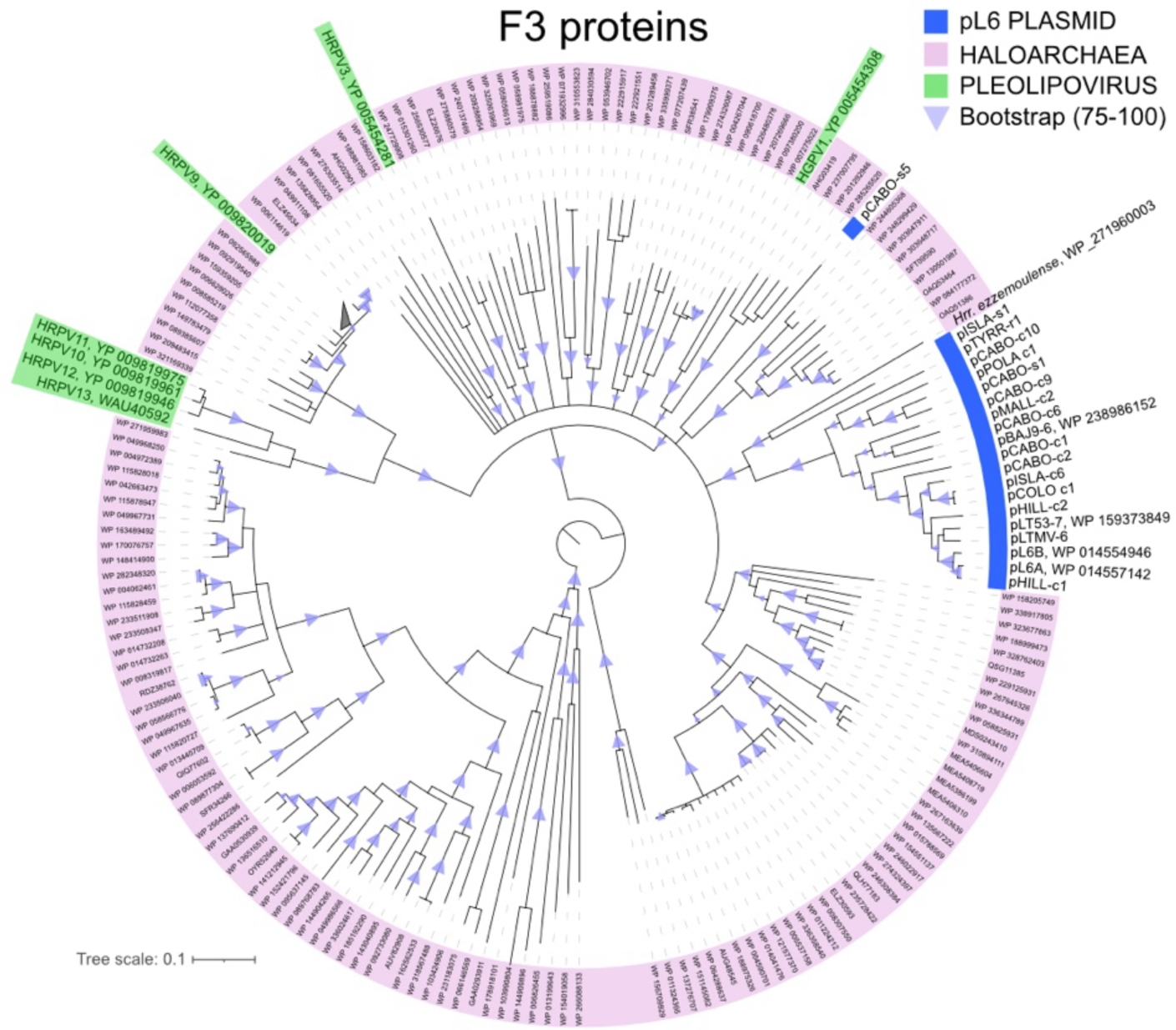
Inferred phylogeny of pL6-family F3 (replicase) proteins. Neighbor-joining consensus tree from 1000 bootstrap repetitions (see Methods). Branches with significant bootstrap confidence values are indicated by light purple triangles. Nodes with accessions in pink shading represent proteins from members of the class *Halobacteria*. Thick blue underlining, pL6-family plasmids. Light green shading, pleolipoviruses (both accession and virus abbreviation are shown, e.g. HRPV3, Halorubrum pleomorphic virus 3).

**Figure 6.**
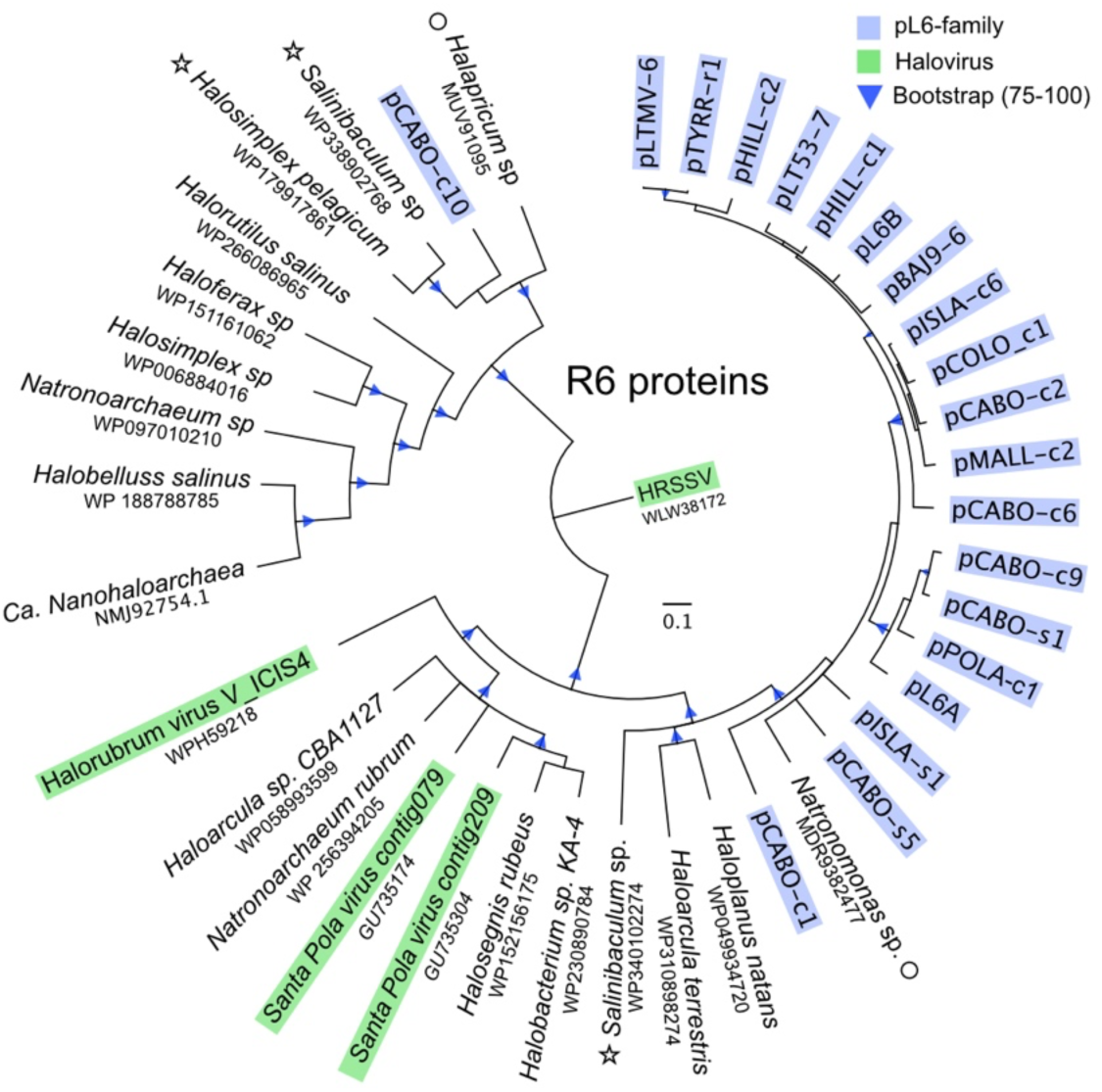
Inferred phylogeny of pL6-family R6 (ATPase) proteins. See legend to previous figure for details of tree and general labelling. A circle next to the species name indicates the protein sequence comes from a small DNA contig, and an asterisk indicates the gene is part of a plasmid-like element (see Text).

In the F3 tree (Figure 5), only the proteins of the two high GC plasmids (pCABO-s5 and pISLA-s1) branch separately from the other pL6-family members. Both show long branch lengths and no close relationship to known haloviruses or to proteins with a gene neighborhood that provided any informative pattern.

In the R6 tree (Figure 6), the pCABO-c10 protein branches separately from the other pL6-family proteins and forms a clade with cellular homologs of *Salinibaculum* (WP338902768) and *Halosimplex* (WP179917861, HZS54_14695 in QLH82792.1). The genes for both proteins are chromosomal and their gene contexts show they are part of integrated elements of about 6.5 kb (Figure S9). These elements share 59.6% nt identity and are flanked at one end by a tRNA-Ala gene and at the other by an integrase gene followed by a short direct repeat matching the end of the tRNA. They share similar gene arrangements and encoded proteins, including an alphapleolipovirus type replicase (see legend to Figure S9), but no genes for virus spike proteins. These two, rare genetic elements appear to represent a group of integrative plasmids possibly related to alphapleolipoviruses, and apart from the integrase gene they superficially resemble the size and gene organization of pL6-family plasmids.

At the other side of the R6 tree, three of the four closest relatives of pL6-family R6 proteins are either encoded on very short contigs or their gene contexts do not provide any obvious clues, but the *Salinibaculum* protein WP340102274 is found on a 5,984 bp contig with directly repeating ends that probably represents a small, circular plasmid. It carries a gene for a putative replicase (WP340102282, DUF1424 domain) that is unrelated to the F3 replicase of pL6-family plasmids or haloviruses.

### 3.5. Conserved DNA sequence motifs

Promoters and regulatory sequences that drive and regulate expression of the outwardly facing gene modules of pL6-family plasmids are probably located in the intergenic region between the R4 and F1 genes (Figure 1), upstream to both. Two conserved intergenic sequences (CIS) in this region were described previously, one of which (CIS 1) contained a typical haloarchaeal promoter motif and the other (CIS 2) possibly being regulatory [13]. In the expanded set of twenty plasmids the two CIS regions were conserved in 17 cases (Figure 7). The three variants not included in the alignment are pCABO-s1 and the two high GC plasmids (pCABO-s5, pISLA-s1).

**Figure 7.**
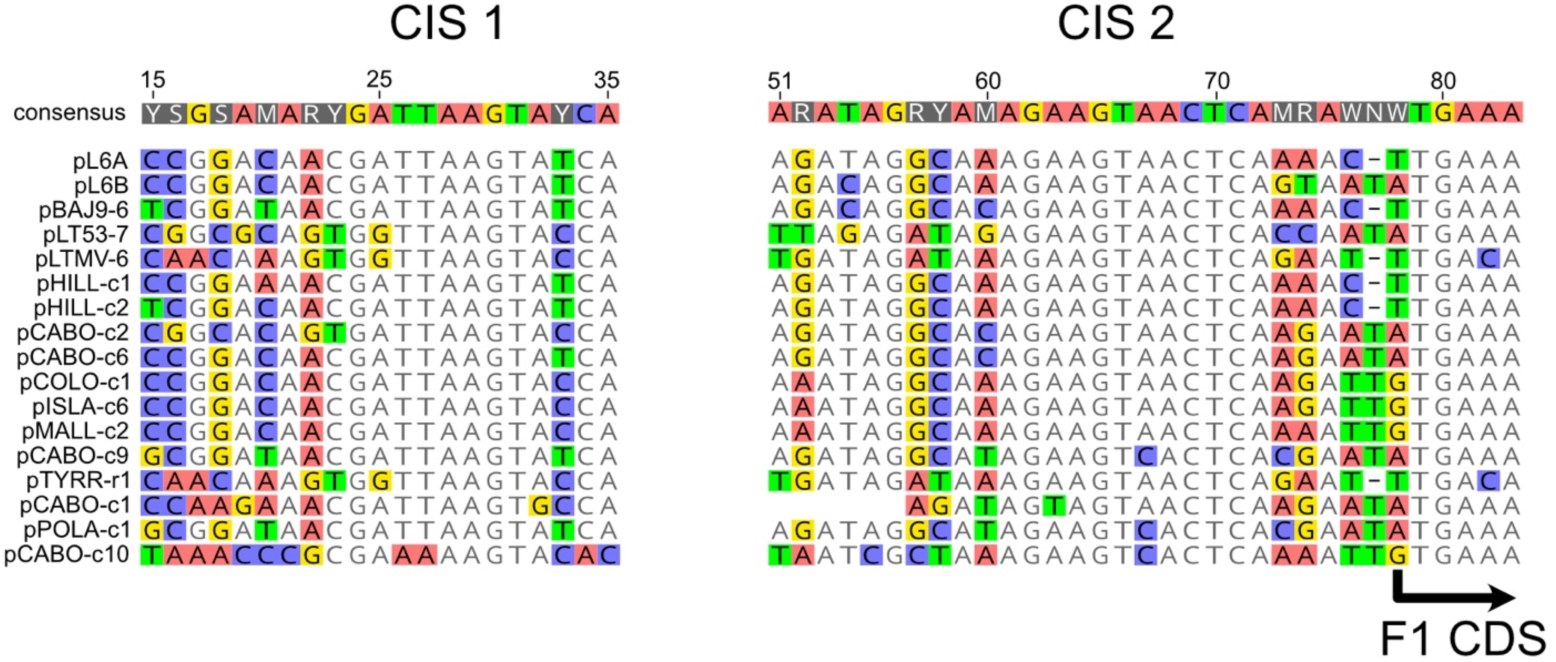
Conserved intergenic sequences (CIS) found between genes R4 and F1. Plasmid names are given at the left, and nucleotide numbers for pL6A are shown at top. Bases are colored if they differ from the consensus at that position (threshold, 75%). The start codon for gene F1 is indicated by the arrow at lower right.

### 3.6. CRISPR spacers that match reconstructed pL6-family plasmids

A total of 109 spacers showing close similarity to one or more of the 15 plasmids reconstructed in this study were retrieved from the IMG/JGI CRISPR spacer database or from metagenomes using the minCED tool (see Methods) and are summarized in Table S5. All came from metagenomes of hypersaline lakes or saltern crystallizer ponds, including the metagenomes used for plasmid reconstructions. Many matching spacers came from sites that were geographically distant to the sites from which the plasmids were derived, and this is depicted in Figure 8 for two plasmids, pCABO-c2 (Puerto Rico) and pCOLO-c1 (Argentina). Matching sequence positions are indicated above each gene map, along with the country of origin of each spacer. Matching sites are almost always located within protein coding sequences, particularly the F3 gene encoding the repli-case.

**Figure 8.**
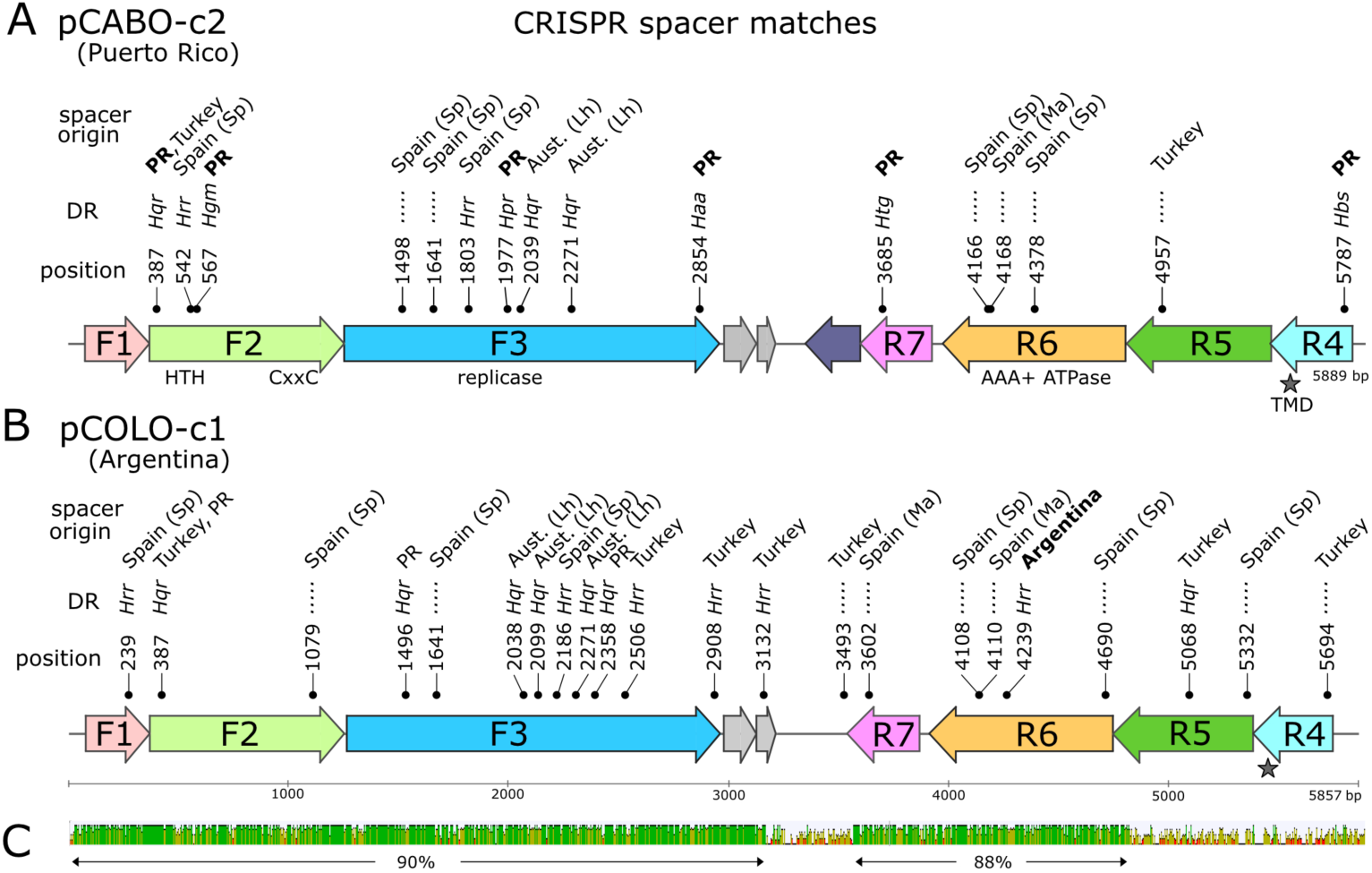
Gene map of plasmids pCABO-c2 (A) and pCOLO-c1 (B) showing positions of matching CRISPR spacers. For gene labeling see legend to Figure 1, and for spacer details see Table S4). Positions of matching spacers are indicated above the map with labels indicating the first matching base (e.g. 387). DR (direct repeat) level above indicates the genus (3-letter abbreviation) of the organism with the most closely related direct repeats associated with each spacer. Haa, *Halanaeroar-chaeum*; Hbs, *Halobellus*; Hgm, *Halogeometricum*; Hpr, *Halapricum*; Hqr, *Haloquadratum*; Hrr, *Halorubrum*; Htg, *Haloterrigena*. Dotted lines indicate no DR match was detected. Top level indicates the country of origin for each matching spacer: PR (Puerto Rico); Turk., Turkey; SP, Spain (Sp, Santa Pola; Ma, Mallorca); Aust. (Lh), Australia (Lake Hillier). Bold face indi-cates that the plasmid and the spacer are from the same country. C, nucleotide similarity of the two aligned plasmids (MUSCLE aligner) with regions of identity represented as green bars. Two regions of high similarity are indicated by arrows below along with their average nucleotide identity values.

The direct repeats (DR) flanking these spacers could be used to identify the genera carrying these CRISPR arrays, and so the likely hosts of pL6 plasmids (see Methods). Over 60% of DR (66/109) could be matched to known genera, and all of them were haloarchaea (Table S5). Half of these (33) matched DR of *Haloquadratum* and were distributed across 10 of the 15 plasmids. In the case of pCABO-c1, all eight of the spacer-associated DR matched those found in *Haloquadratum*. For pHILL-c2, five DR out of eight could be matched and all five corresponded to DR of *Haloquadratum*. The likely hosts for both pCABO-c1 and pHILL-c2 are members of the genus *Haloquadratum*, and this is also supported by the accessory genes they carry (Table S4). The remaining 33 DR matched a total of ten other genera of the class *Halobacteria*, most of which are commonly detected in salt lakes and crystallizer ponds, such as *Halorubrum* and *Halogeometricum* [8]. Overall, the evidence from CRISPR spacers and DRs is consistent with pL6-family plasmids being present in microbial communities of hypersaline environments around the world, and with their frequent invasion of haloarchaea, particularly *Haloquadratum*.

### 3.7. RNA-seq evidence for transcription of pL6-family plasmid genes

RNA-seq data of the 2018 microbial community of Lake Tyrrell (SRR24125903) is available from the study of [10]. When these reads were mapped to all pL6-family plasmids using the Geneious Prime ‘map to reference’ tool (threshold of ≤2% mismatches, see Methods) by far the most reads matched to pLTMV-6 (367 reads), a plasmid that was reconstructed from a Lake Tyrrell metagenome and described previously [13]. In order to check if more closely matching plasmids to the RNA-seq reads could be recovered from Lake Tyrrell metagenomes, the RNA-seq reads mapping at low stringency (up to 35% sequence variation) to the F3 gene of pLTMV-6 were de novo assembled, and the contig with the highest number of reads used as a seed sequence to extend at both ends by repeated read mapping with 2009 Lake Tyrrell metagenomic (DNA) reads (SRR5637210-11). This resulted in a 5,844 bp plasmid designated pTYRR-r1 (Table 1) that is closely similar to pLTMV-6 in genes F1 and R4-R7 but has lost the F2 gene and differs significantly in sequence at the 3’ end of the F3 gene (Figure S10).

A total of 385 RNA-seq reads matched the pTYRR-r1 sequence when a more stringent threshold of ≤1% base mismatches was applied (Figure 9, upper panel). All but 38 nt of the plasmid was represented by RNA-seq reads, with a distinct peak of read coverage (24.5 x) across the gene for an insertion sequence (ISH) and an adjacent CDS (nt 2379-3107). Since the RNA-seq data of Le Lay et al. is strand specific (Illumina TruSeq), the reads have been color-coded in the figure to indicate the strand (grey, top strand; pink, lower strand).

**Figure 9.**
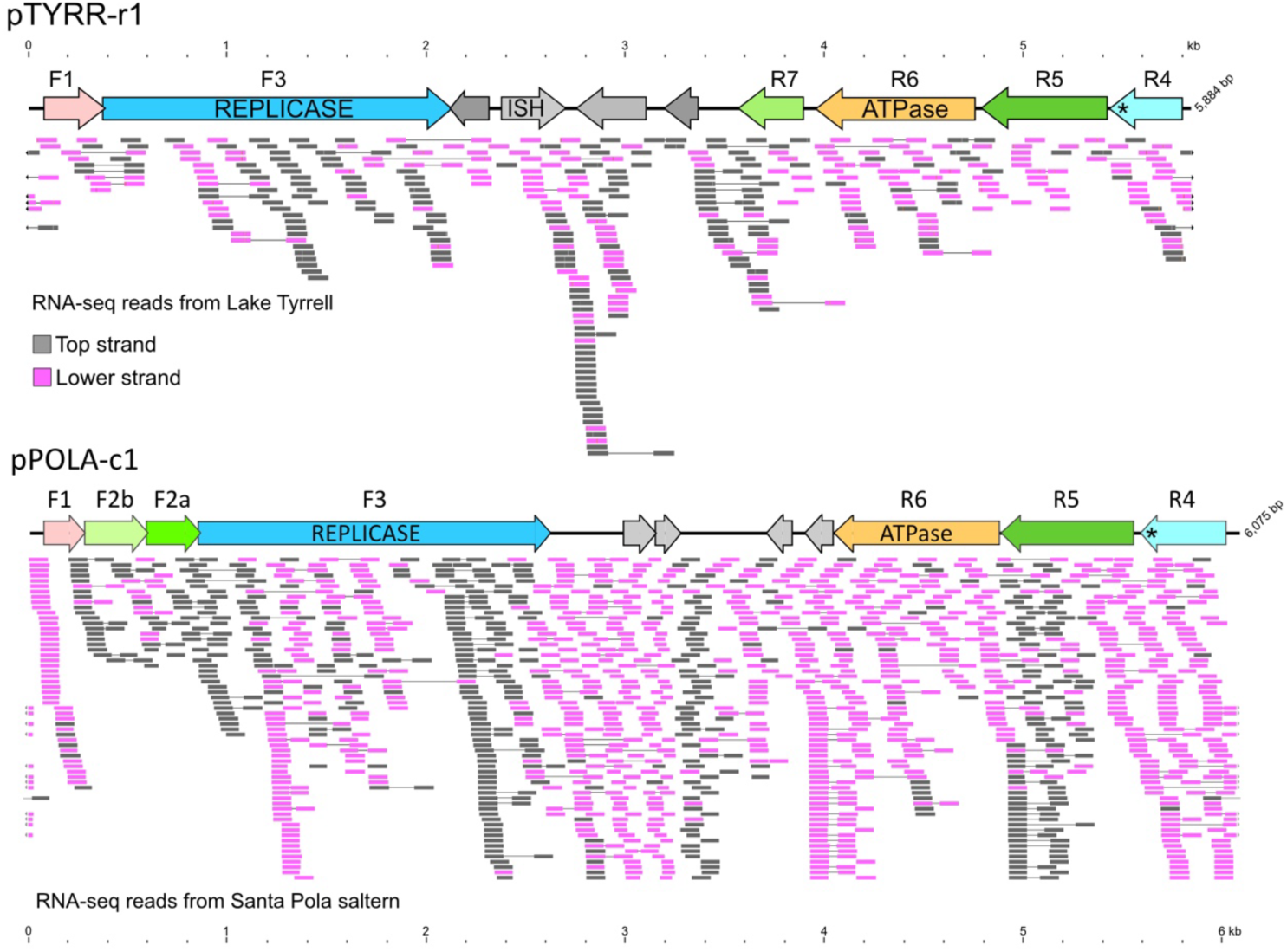
Metatranscriptome reads from Lake Tyrrell (SRR24125903) and Santa Pola saltern (SRR10674835-40) were mapped to the sequences of plasmid pTYRR-r1 (upper panel) and pPOLA-c1 (lower panel), respectively. Mapping was done within the Geneious Prime environment (see Methods) with maximum base mismatch threshold of 1%. Mapped reads are represented below each gene map and are color-coded according to strand: grey (top strand); pink (lower strand). Lines connecting reads indicate well separated read pairs. Arrows at edges represent reads spanning the two edges. A scale is given above the pTYRR-r1 gene map in the upper panel, and below the mapped reads of the lower panel. For clarity, most of the reads mapping to the middle of pPOLA-c1 gene map, where the read coverage is extremely high, have been removed, leaving an average coverage = 23.

Metagenomic RNA-seq data is also available for the Santa Pola saltern (SRR10674835-40) and these were mapped to pPOLA-c1, a plasmid reconstructed in the present study from the metagenome of the same saltern (Figure 9, lower panel). A total of 78,582 paired reads mapped to this plasmid (stringency of ≤ 1% mismatches) but most reads matched two adjacent, small CDS located near the middle of the plasmid (280 bp; nt 2992-3271) in a region where insertions of foreign genes are often found in other pL6-family plasmids [13]. These two small CDS are similar to chromosomal sequences of *Haloquadratum walsbyi* that often occur nearby HqIRS55 insertion sequences (e.g. FR746099.1, nt 1362480-1362865). Ignoring the central, high-coverage region, reads matching upper and lower strands were common throughout the entire sequence of pPOLA-c1.

The presence of reads mapping to both strands across most of the annotated genes of both plasmids indicates not only active transcription of their genes but also widespread counter-transcription. High levels of counter-transcription have been observed in haloviruses such as HF2 [35] and SH1 [36].

## 4. Discussion

Fifteen pL6-family plasmids were reconstructed in this study from the metagenomes of salt lakes and saltern crystallizer pools of four widely separated countries: Argentina, Australia, Puerto Rico and Spain. This increases the number of complete plasmid sequences from five to twenty and greatly expands their geographical distribution. All share the archetypical gene arrangement of outward facing replication and ATPase modules reported previously [13], and while nearly half of the plasmids (7 of 15) displayed a closely similar gene complement to the five earlier examples, some novel variants were discovered, such as those where the F2 gene has been replaced with two smaller genes (F2b/F2a) or lost entirely. The significance of these differences remains unresolved as the molecular functions of the proteins encoded on pL6-family plasmids are largely unknown, and their sequences provide few clues.

The association of pL6-family plasmids with *Hqr. walsbyi* seen previously [13] is supported by the findings of the current metagenomic study. CRISPR DR sequences indicated that at least two plasmids (pCABO-c1, pHILL-c2) are from this species. This is also consistent with the accessory (acquired) genes of these two plasmids, which have encoded proteins most similar to homologs from *Hqr. walsbyi*. The high community abundance of *Haloquadratum* in the Puerto Rico (53.8 to 69.8%), Lake Tyrrell (58%) and Santa Pola (63%) saltern ponds [1,8] also makes it likely that plasmids recovered from these sites are harbored by this species, which would include the pCABO-series (except pCABO-s5, see below), pTYRR-r1 and pPOLA-c1. In the cases of pCABO-s1 and pTYRR-r1, their accessory genes also support a *Hqr. walsbyi* host. Although DR sequences from genera other than *Haloquadratum* were observed, suggesting pL6-family plasmids can spread to other species, no examples have yet been reported in the literature, nor can pL6-family plasmids be found in the currently available (July, 2024) complete genome sequences of cultivated haloarchaea. Why these plasmids appear to be most commonly carried by *Hqr. walsbyi* remains to be unraveled.

The avoidance of specific palindromic motifs is common in mobile genetic elements and viruses of prokaryotes in order to circumvent restriction-modification (RM) defense systems of host cells [37,38]. Many RM systems of haloarchaea have been documented [39,40]. In the current study, tetramer analysis confirmed the general avoidance of GGCC and CTAG motifs in pL6-family plasmids as was reported previously [4], but in this larger dataset there were two prominent exceptions. The two high GC plasmids pCABO-s5 and pISLA-s1 maintain both motifs at the expected frequency. Since the sequenced strains of *Hqr. walsbyi* (C23T and HBSQ001) avoid GGCC and CTAG motifs [4], and carry multiple restriction systems, one of which is predicted to recognize GGCC motifs (Hqrw_2139; REBASE, http://rebase.neb.com, accessed June 10, 2024), the atypical sequences of pCABO-s5 and pISLA-s1 suggest their usual hosts are haloarchaeal species other than *Haloquadratum*.

The case of pHILL-c2, where GGCC is absent and CTAG sites are not under-represented is best explained by the acquisition of foreign DNA by this plasmid that contains multiple CTAG sites. This acquired DNA carries an accessory gene encoding a formyltransferase, and using the predicted 3D structure of this enzyme it could be identified as a sugar N-formyltransferase that probably synthesizes dTDP-4-formamido-4,6-dideoxyglucose. In bacteria like *Mycobacterium tuberculosis*, the N-formylated sugar ends up in cell surface polysaccharides [33,34]. Further evidence for its natural function was sought by examining the gene neighborhoods for closely matching proteins, which showed adjacent genes commonly encoded glycosyltransferases or enzymes involved in polysaccharide biosynthesis, supporting the function obtained from 3D structure similarity results. Two other accessory genes carried by pL6-family plasmids were examined in the same way. Both coded for methyltransferases, and the genes surrounding the most closely matching proteins were frequently glycosyltransferases. Taken together, these results suggest that enzymes involved in sugar modification or addition provide a selective advantage for pL6-family plasmids, possibly involving alteration of glycans attached to cell surface proteins or cell surface polysaccharides.

The possibility that pL6-family plasmids represent a novel group of temperate viruses was suggested by the studies of Lake Tyrrell, where water filtered down to 0.1 µm was used to produce ‘virus concentrates’ [41], the DNA of which had abundant reads to this plasmid family and were used to reconstruct plasmid pLTMV-6. However, filtration not only recovers virus particles but also dissolved DNA released from lysed cells, extracellular membrane vesicles (EV)[42], and membrane enveloped plasmids [43,44]. Viruses are normally purified by a combination of methods that usually include density gradient centrifugation and ultracentrifugation, and in the recent study of [11] the authors fractionated Santa Pola crystallizer water into dissolved DNA (by filtration) and virus pellets (by ultracentrifugation). The DNA sequence reads from those fractions were used in the present study to determine the relative abundance of plasmid pPOLA-c1 in these fractions. Compared to cellular DNA, pPOLA-c1 reads were 1.6-2.6x higher in dissolved DNA fractions but 34-fold less in the virus pellet fraction. An unexplained over-representation of plasmids in the dissolved DNA fractions was also noted by Aldeguer-Riquelme *et al.,* who found a two-fold higher abundance of *Hqr. walsbyi* plasmid pL47 in dissolved compared to intracellular DNA [11], and they speculated that this might be due to plasmid resistance to exonucleases, or by export from cells in membrane vesicles [44]. In any case, the very low levels of pPOLA-c1 in the virus pellet is evidence against it forming virus particles that could be sedimented by ultra-centrifugation. Three possible interpretations of this data are, that (a) pPOLA-c1 exists only as a plasmid, (b) pPOLA-c1 is a temperate virus but induction rates are extremely low, and (c) the fractionation methods used by [11] did not pellet low-density or delicate (e.g. lipid enveloped) viruses. Indeed, their analysis identified only tailed viruses (Class *Caudoviricetes*) in virus pellets.

Transcription of pL6-family plasmids has not been reported previously and was only possible because of publicly available strand-specific metatranscriptome data from Lake Tyrrell and Santa Pola that could be interrogated using plasmids reconstructed from metagenomes obtained from the same sites. The results provide unexpected insights into plasmid gene expression. Hundreds of closely matching, paired reads matched sequences on both strands and across all genes of pTYRR-r1 and pPOLA-c1. The matching transcriptome reads provide independent support for the reconstructed plasmid sequence of pPOLA-c1, and for most of pTYRR-r1. While there are many unknown variables in these results, they imply (a) significant levels of these plasmids were present in the corresponding microbial communities, sufficient for the detection of their transcripts, and (b) that widespread transcription occurs from both DNA strands, producing both sense and antisense transcripts. Antisense or counter-transcripts have commonly been observed in transcriptome studies of prokaryotes [45], including haloarchaea [46–49], and are generally thought to be involved in regulation of gene expression [46]. A study of *Halobacterium salinarum* found that at least 21% of genes produced antisense transcripts but most were at low levels [48]. In the haloviruses that have been studied, antisense transcripts are commonly detected [36].

Although the additional examples of pL6-family plasmids recovered in this study have shed light on their conserved and variable features, the underlying biology of these plasmids remains poorly understood. CRISPR spacer matches show they invade haloarchaeal cells but how are they transferred or gain entry, and what genes are involved in maintenance or partition? Are they merely plasmids, or defective remnants of temperate pleolipoviruses, or perhaps a novel type of virus? Their sequences show features similar to lipid enveloped pleolipoviruses [15,50,51], particularly the betapleolipovirus group which have a similar replicase gene and, like many other viruses, display a strong conservation of gene arrangement with colinear, often overlapping genes that form functional modules. Such arrangements allow viruses to rapidly and efficiently produce virions during productive infection while keeping genome sizes within packaging limits. In contrast to viruses, plasmids have far fewer constraints and are more likely to vary in size and gene complement [52,53]. While pL6-family plasmids maintain a conserved gene arrangement and a limited size range they do not encode a membrane anchored protein related to the spike proteins of pleolipoviruses (or any other virus), yet the capsid proteins of viruses are generally well conserved in structure, permitting relationships to be discerned across wide evolutionary distances [54]. The only predicted membrane anchored protein encoded by pL6-family plasmids is R4. Finally, ultracentrifugation of filtered saltern water from Santa Pola [11] showed no evidence that pL6-family plasmids are packaged within capsids or lipid envelopes. Our analysis provides a foundation for future studies to elucidate the true nature of these intriguing genetic elements, the functions of their proteins, and how they regulate gene expression, including the role of antisense transcripts.

## Supporting information

Supplementary Figure S1, panels A-O

Supplementary Figure S2, panels A-I

Supplementary Figure S8, panels A-C

## Supplementary Materials

See link on BioRxiv download page

## Author Contributions

M.DS.; conceptualization, investigation, formal analysis, writing. F.P.; investigation, writing—review and editing. Both authors have read and agreed to the published version of the manuscript.

## Data Availability Statement

Sequence accessions will be available from Genbank upon acceptance of the manuscript for publication

