## Supplementary Figure S1, panels A-O for "Global distribution and diversity of haloarchaeal pL6-family plasmids"

**Supplementary Figure S1, panels A to O:
Read mapping to reconstructed pL6-family plasmids.**

**Legend to panels A to O.** Figures are based on screenshots of the Geneious Prime assembly window, showing the linearised pL6-family plasmid gene map at the top (with a nucleotide scale above) and the matched reads below. The F3 gene is labeled and coloured red. Reads were mapped using the “map to reference” tool (see methods) and the read coverage is indicated by the blue graph at the very top of each figure. Except for pCABO-c1, pCOLO-c1 and pPOLA-c1, which were set to map reads with zero mismatches (0%), the mapping of reads to the other plasmids used 1% maximum mismatch with the following settings: Geneious mapper (Geneious Prime version 2023.2.1) with custom sensitivity, fine tuning = none, minimum mapping quality = 40, only map paired reads which = both map, maximum mismatches per read = 1%, maximum ambiguity = 1. Reads are coloured by read-pair distance (except for pTYRR-r1 where the reads were not paired and are coloured grey), and vary in colour from orange (close/overlapping) through green to blue (most distant). Lines between reads indicate read pairs. Reads at the extreme left or right edges of the figures have grey arrowheads to indicate reads that span across left and right ends of the linearised plasmid sequence.

**Figure S1, A.** Plasmid pCABO-c1 with metagenomic reads (SRR8816317) mapped at 0% mismatch.

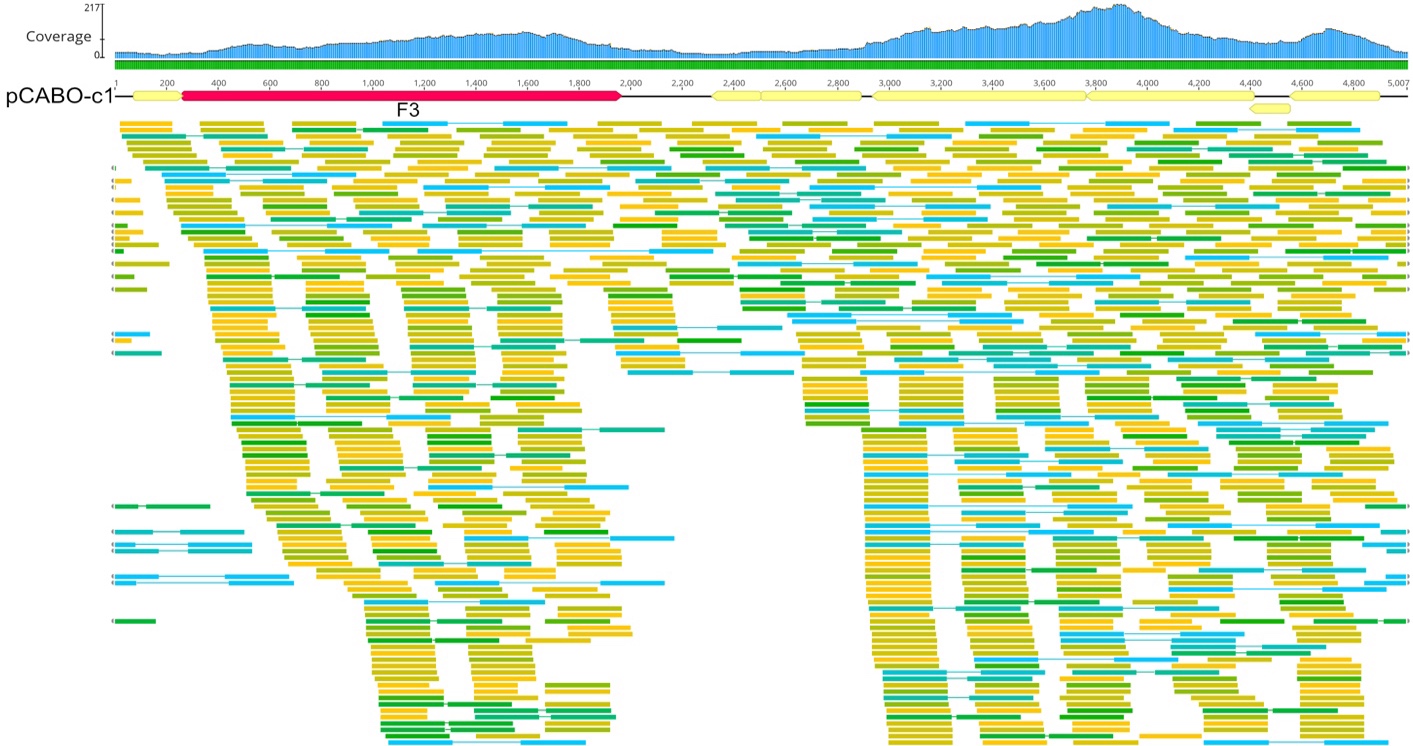

**Figure S1, B.** Plasmid pCABO-c2 with metagenomic reads (SRR8816317) mapped at ≤1% mismatch.

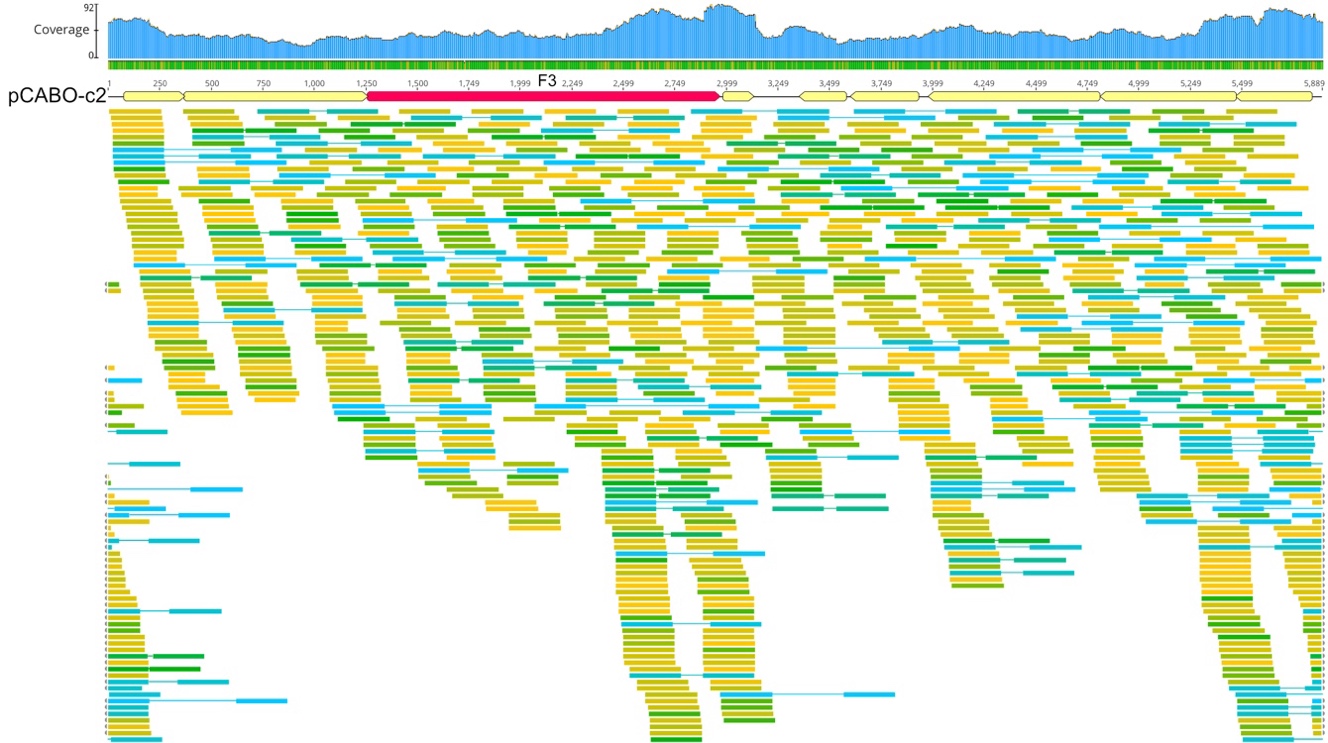

**Figure S1, C**. Plasmid pCABO-c6 with metagenomic reads (SRR8816317) mapped at ≤1% mismatch.

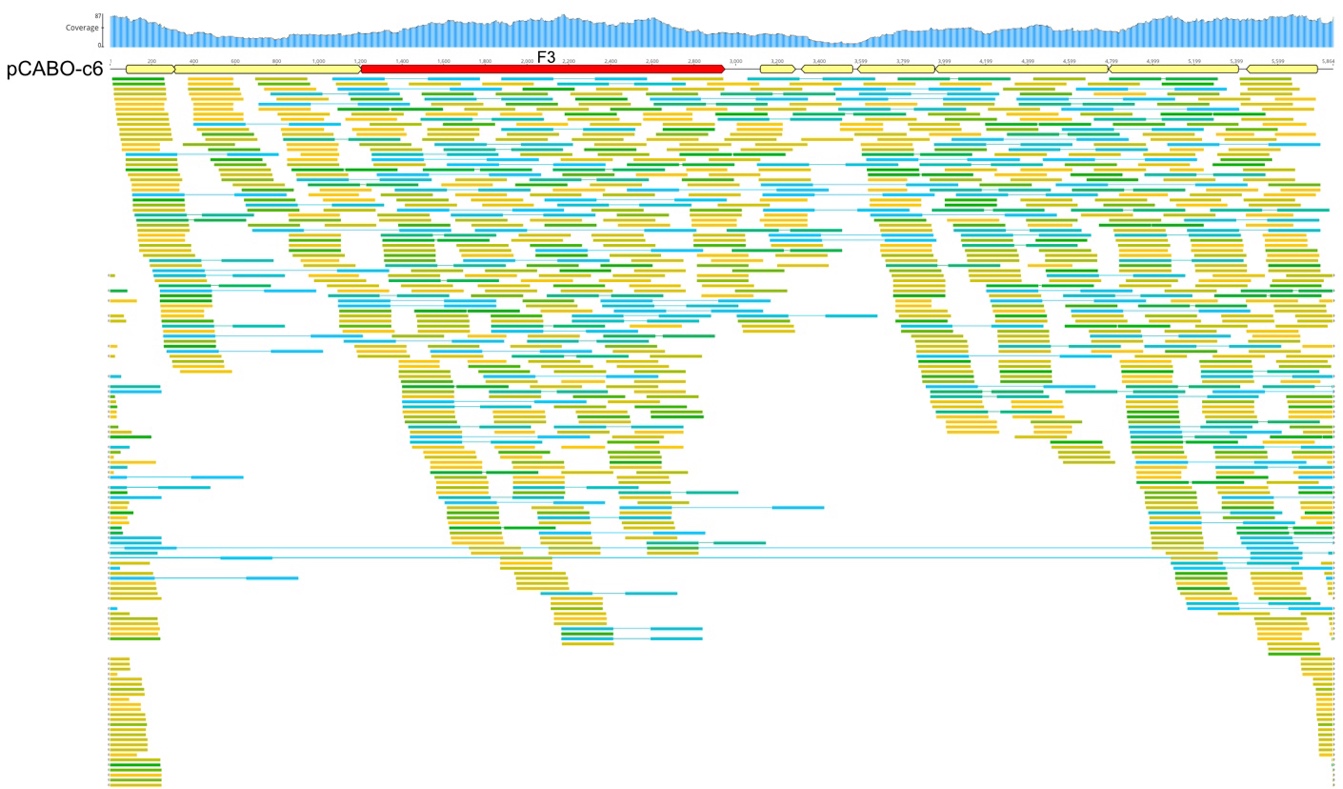

**Figure S1, D**. Plasmid pCABO-c9 with metagenomic reads (SRR8816317) mapped at ≤1% mismatch.

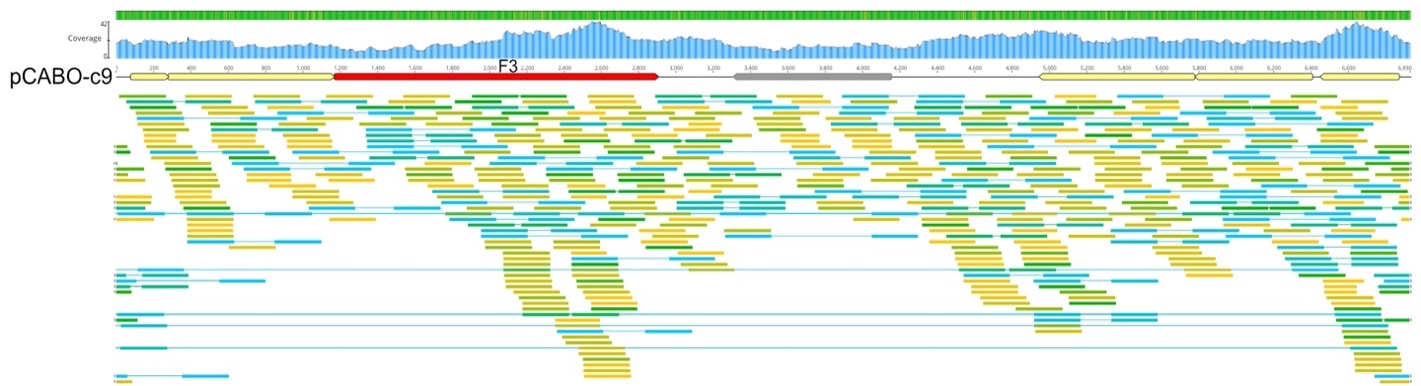

**Figure S1, E.** Plasmid pCABO-c10 with reads (SRR8816317) mapped at ≤1% mismatch.

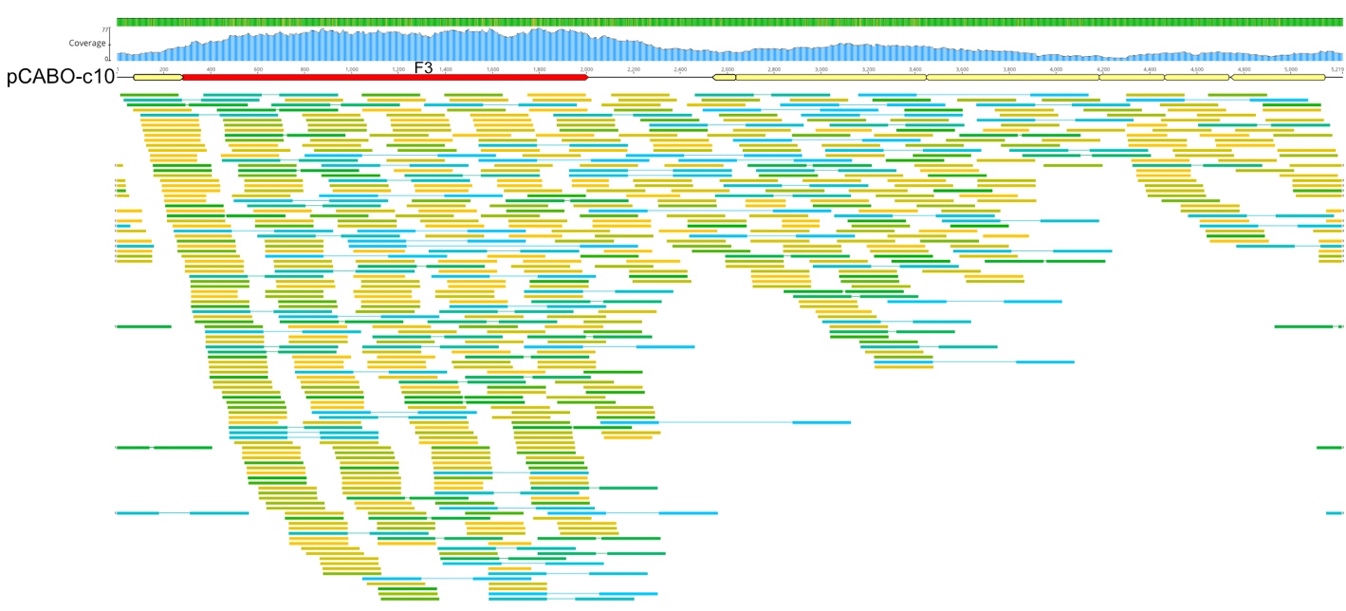

**Figure S1, F.** Plasmid pCABO-s1 metagenomic with reads (SRR8816317) mapped at ≤1% mismatch.

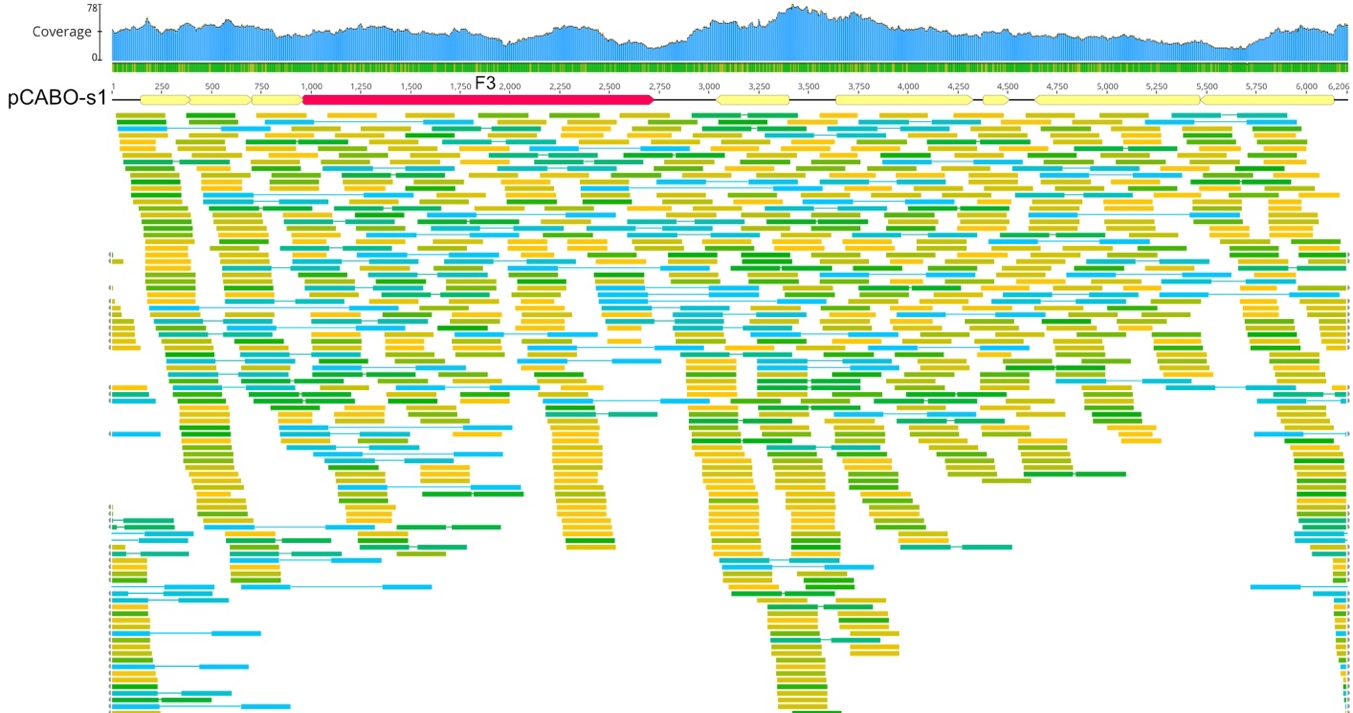

**Figure S1, G**. Plasmid pCABO-s5 with metagenomic reads (SRR8816317-8) mapped at ≤1% mismatch.

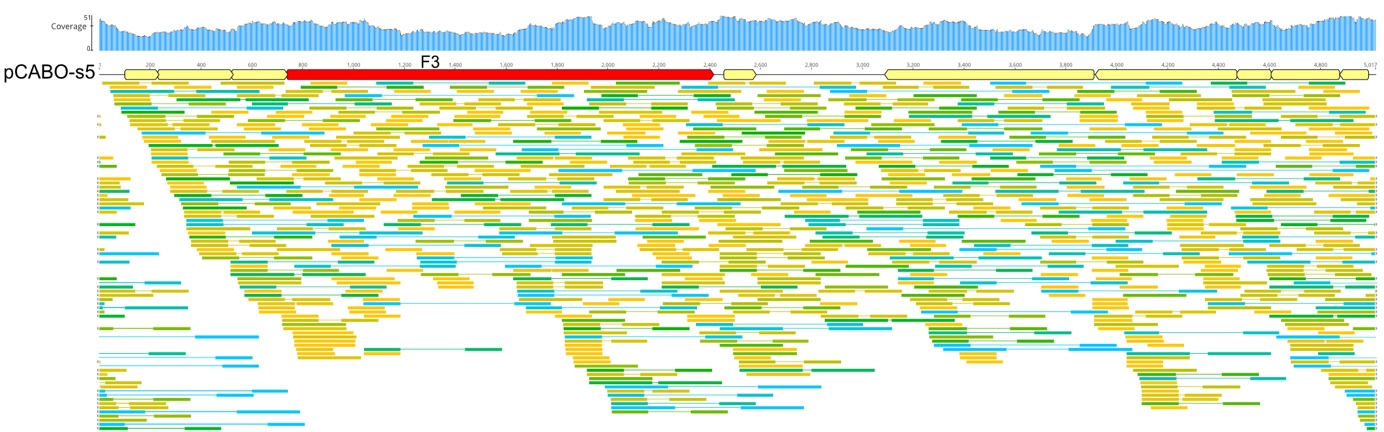

**Figure S1, H.** Plasmid pCOLO-c1 with metagenomic reads (ERR7916263) mapped at 0% mismatch.

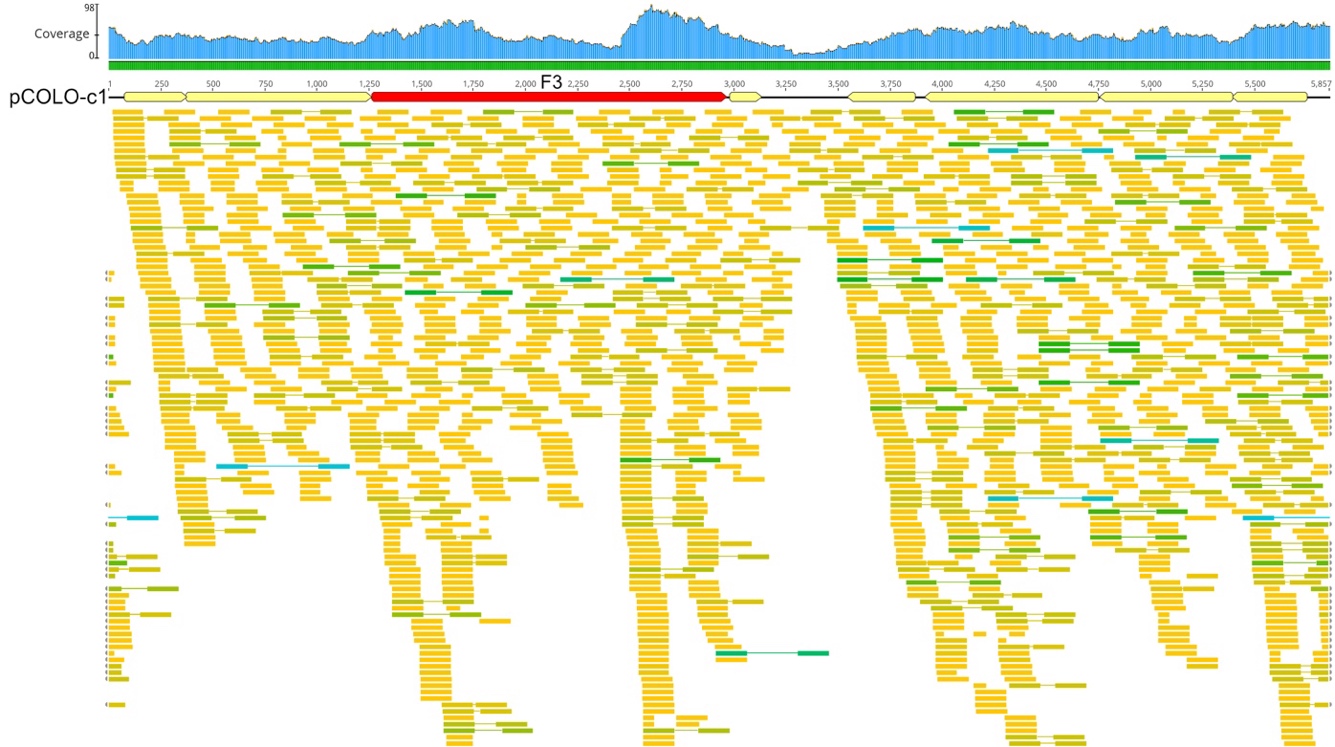

**Figure S1, I**. Plasmid pHILL-c1 with reads (SRR26978067, SRR26968577) mapped at ≤1% mismatch.

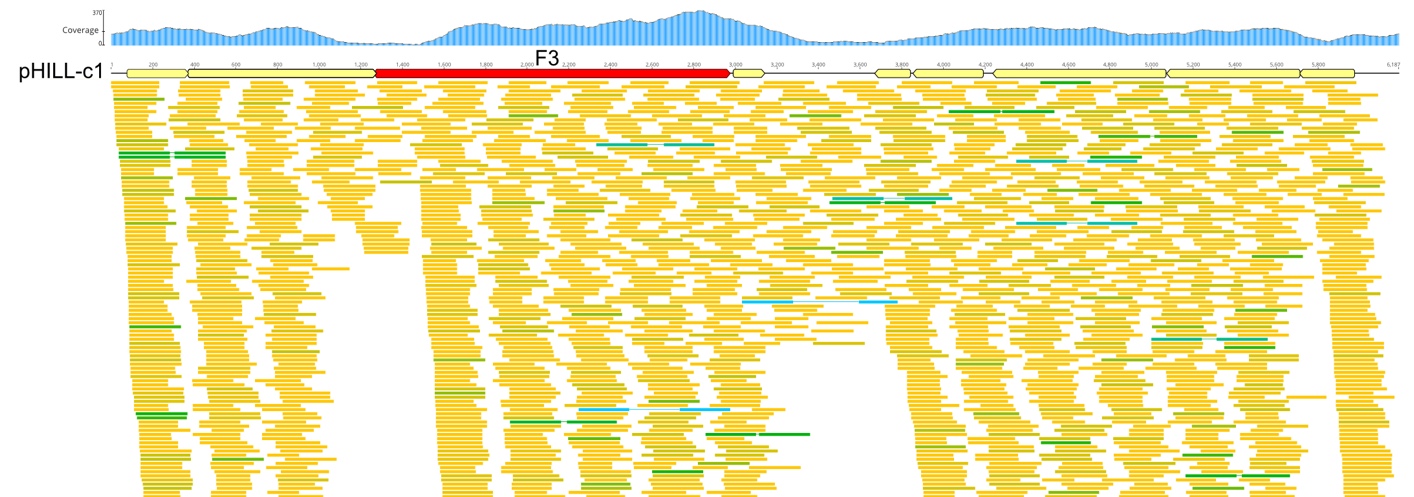

**Figure S1, J.** Plasmid pHILL-c2 with metagenomic reads (SRR26978067) mapped at ≤1% mismatch.

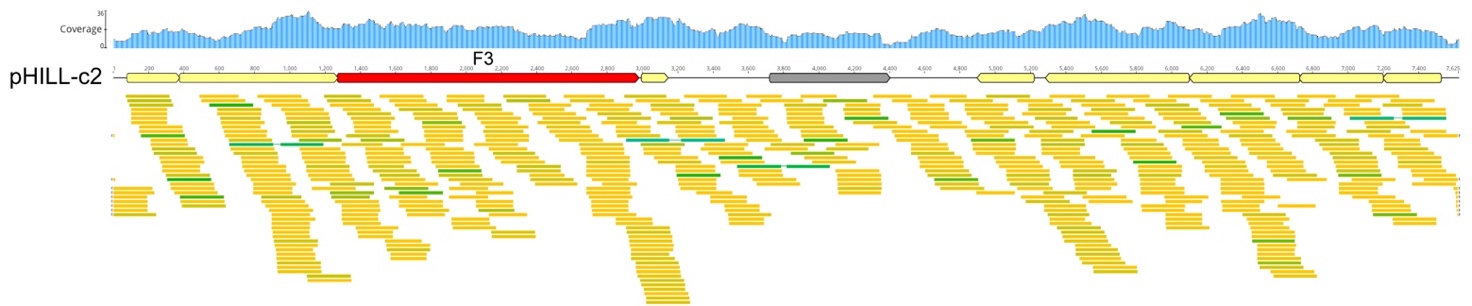

**Figure S1, K.** Plasmid pISLA-c6 with metagenomic reads (SRR21894959, SRR23092357) mapped at ≤1% mismatch.

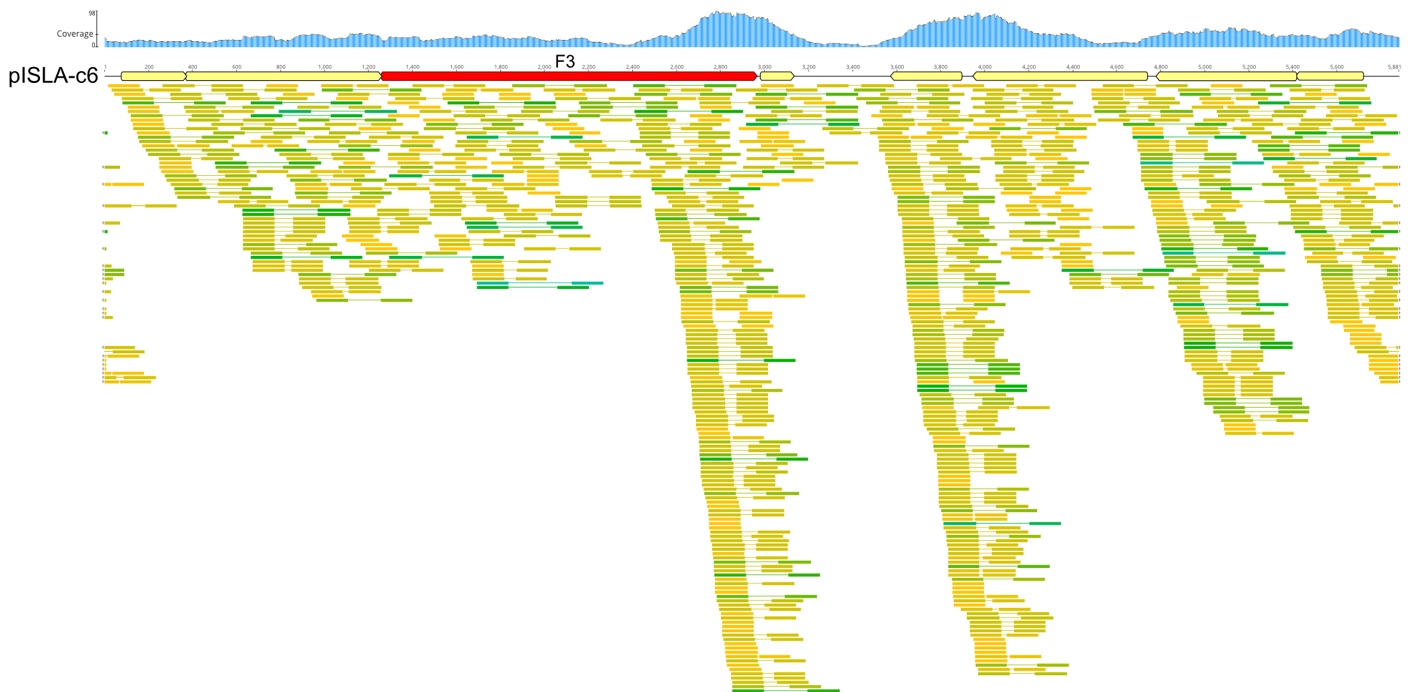

**Figure S1, L**. Plasmid pISLA-s1 with metagenome reads (SRR23092357) mapped at ≤1% mismatch.

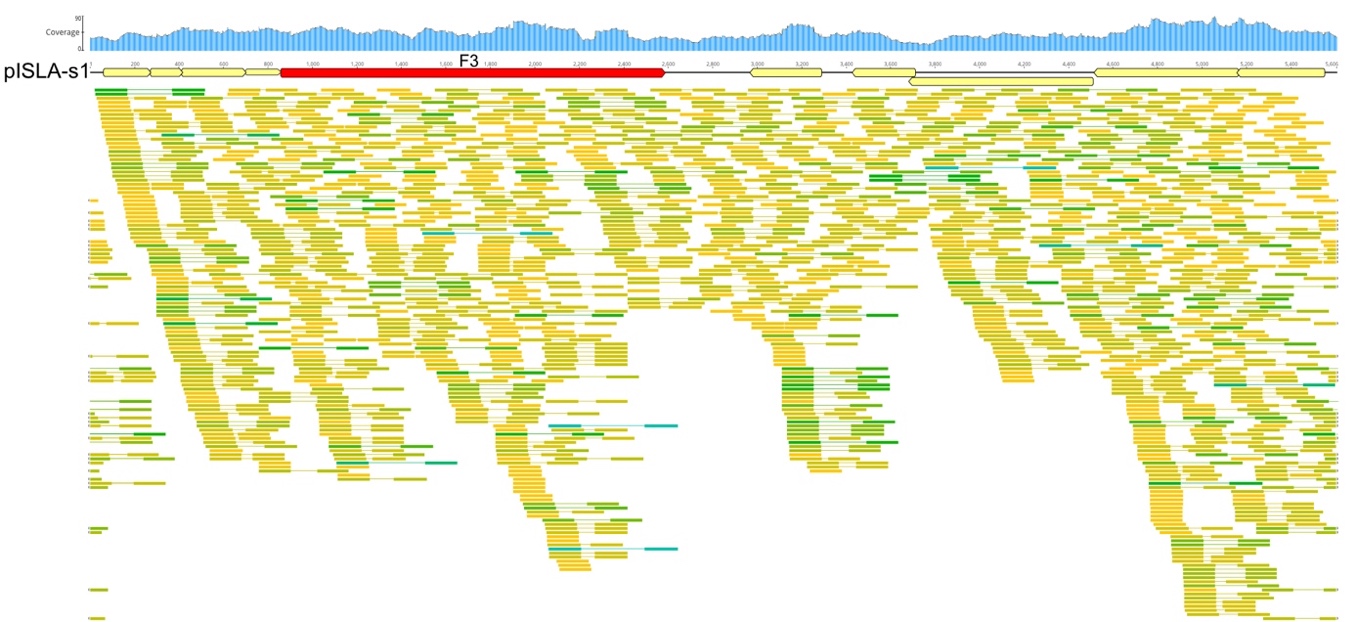

**Figure S1, M.** Plasmid pMALL-c2 with metagenomic reads (ERR5979340) mapped at ≤1% mismatch.

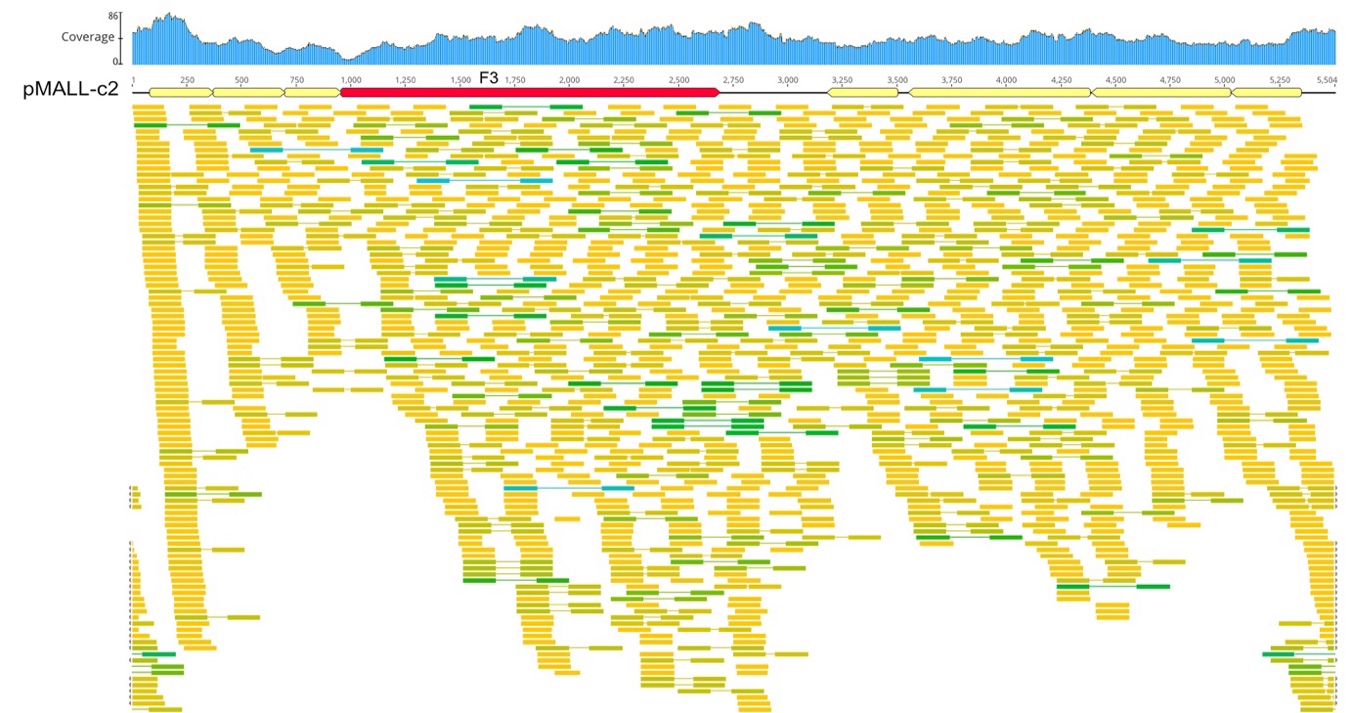

**Figure S1, N.** Plasmid pPOLA-c1 with metagenomic reads (SRR13926770) mapped at 0% mismatch.

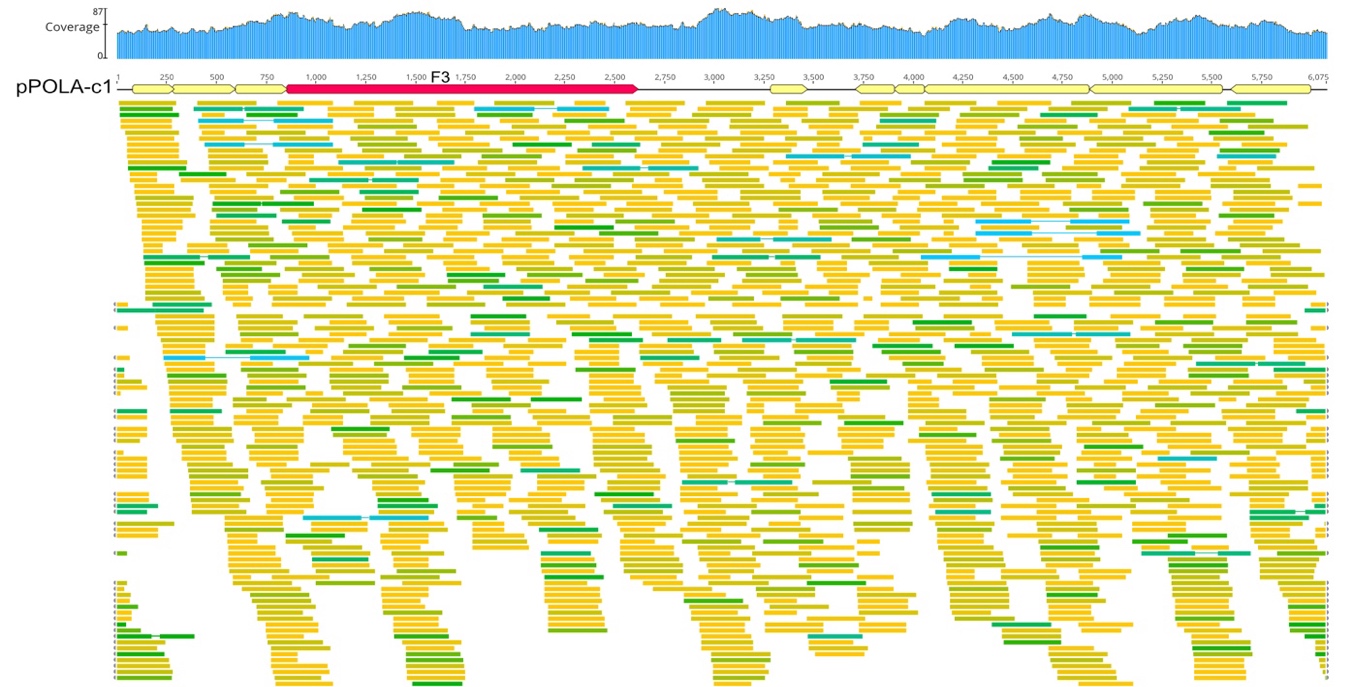

**Figure** **S1, O.** Plasmid pTYRR-r1 with 'virus concentrate' reads (SRR5637210) mapped at ≤1% mismatch.

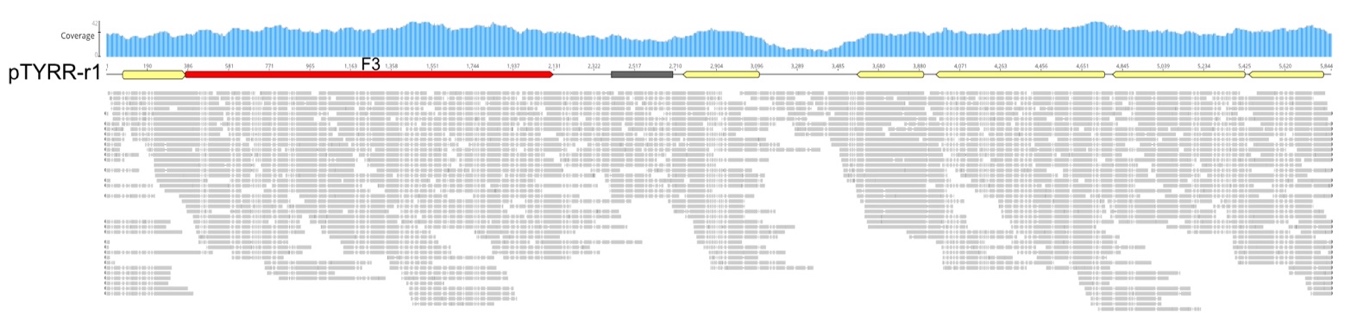
