## Supplementary Figure S2, panels A-I for "Global distribution and diversity of haloarchaeal pL6-family plasmids"

**Supplementary Figures S2A to S2I. Alignments of pL6-family plasmid proteins.**

**Figure S2, A**. Alignment of 18 F1 and two F1b proteins^a^

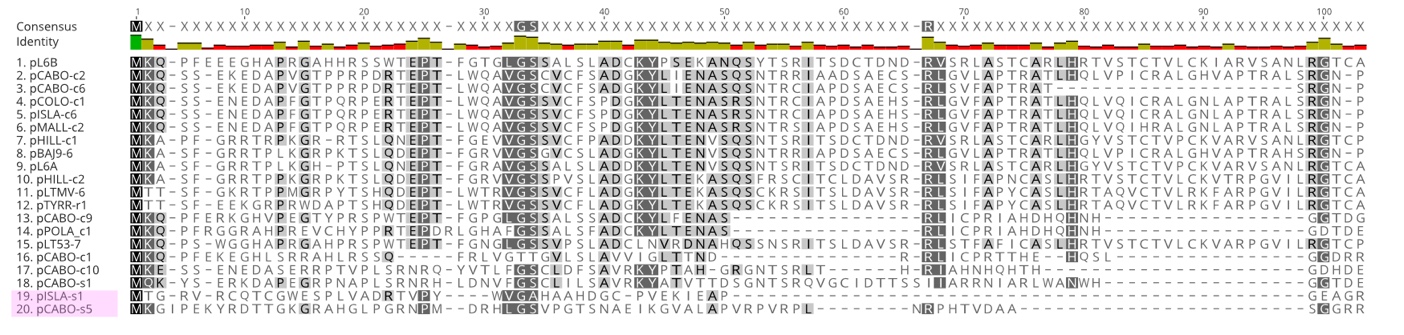

^a^The 18 F1 and two F1b protein sequences from each plasmid (labeled at left) were aligned using the MAFFT v7.490 aligner within Geneious Prime (v.2024.0.5). Residues have been shaded according to similarity, and a consensus identity (threshold of 90%) is shown at the top, along with position numbering. The F1b sequences are indicated by the pink shading of plasmid names.

**Figure S2, B**. Alignment of twelve F2 and five F2a proteins^a^
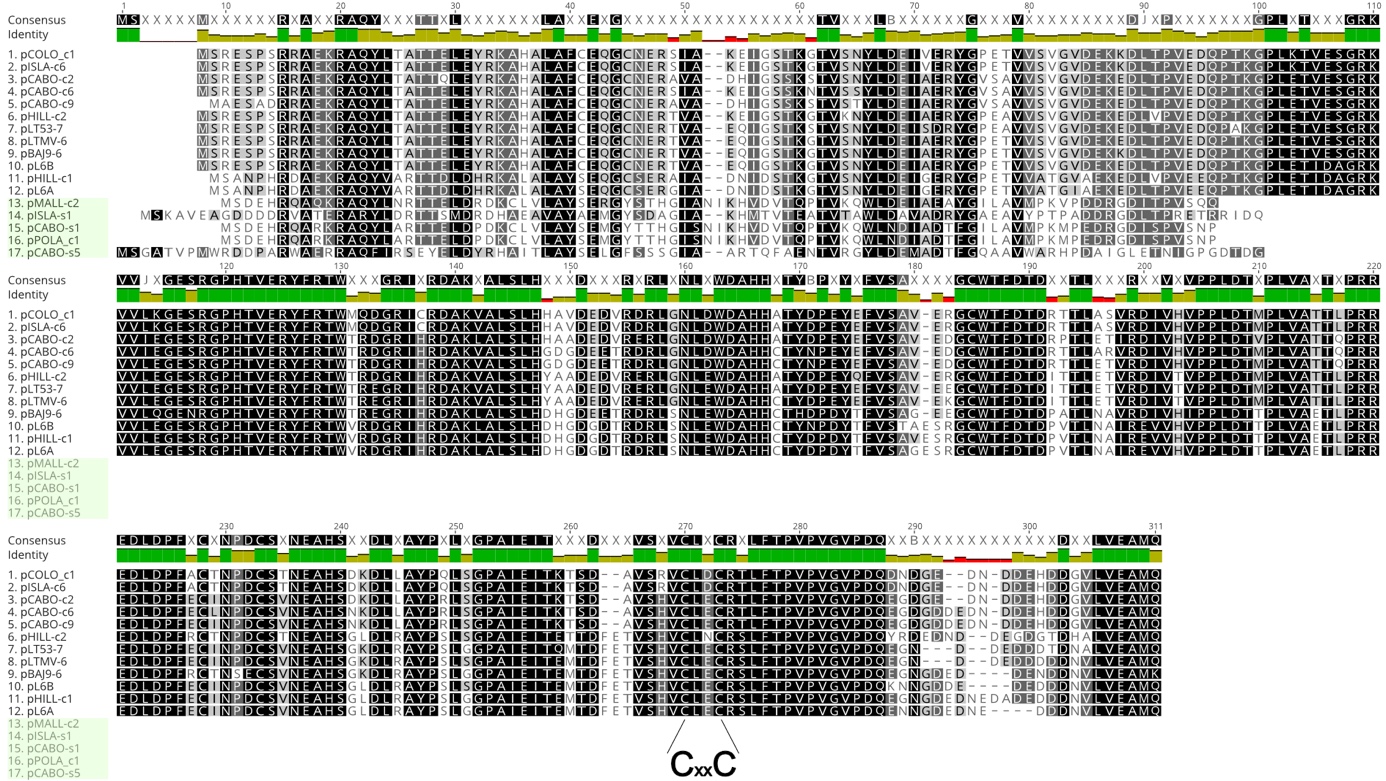

^a^First twelve sequences are F2 proteins, and the remaining five (shorter) sequences are F2a proteins (names highlighted in light green). For other details see legend to Figure S2A (above). Note the CxxC motif beginning at position 270 in the alignment (labeled underneath).

**Figure S2, C**. Alignment of F2b proteins^a^

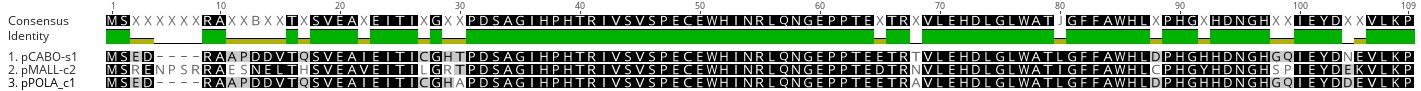

^a^ Inferred F2b protein sequences were aligned according to legend under Figure S2A (above). These proteins do not share similarity with F2, F2a or F2c proteins.

**Figure S2, D.** Alignment of F2c proteins^a^

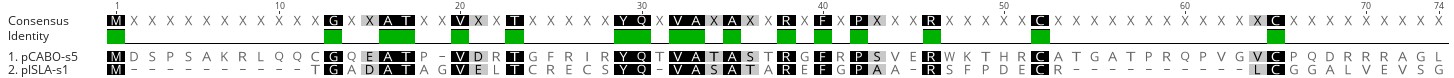

^a^ Inferred F2c protein sequences were aligned according to legend under Figure S2A (above).

**Figure S2, E**. Alignment of F3 (replicase) proteins^a^
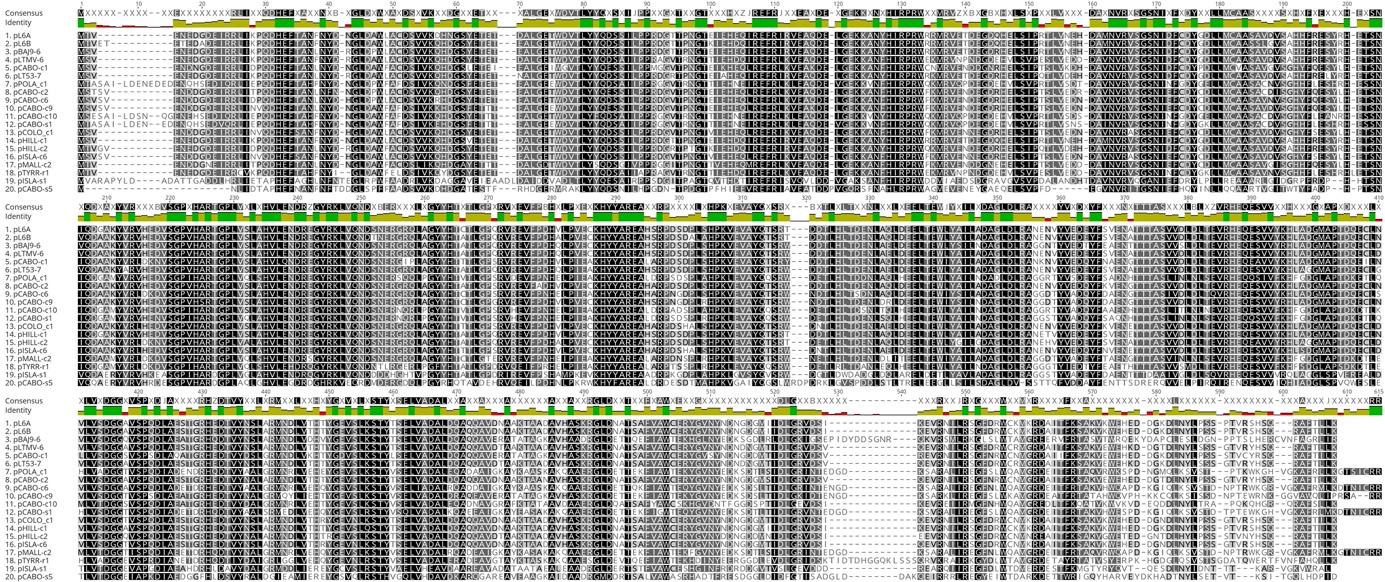

^a^ Inferred F3 protein sequences were aligned according to legend under Figure S2A (above).

**Figure S2, F.** Alignment of R4 proteins^a^
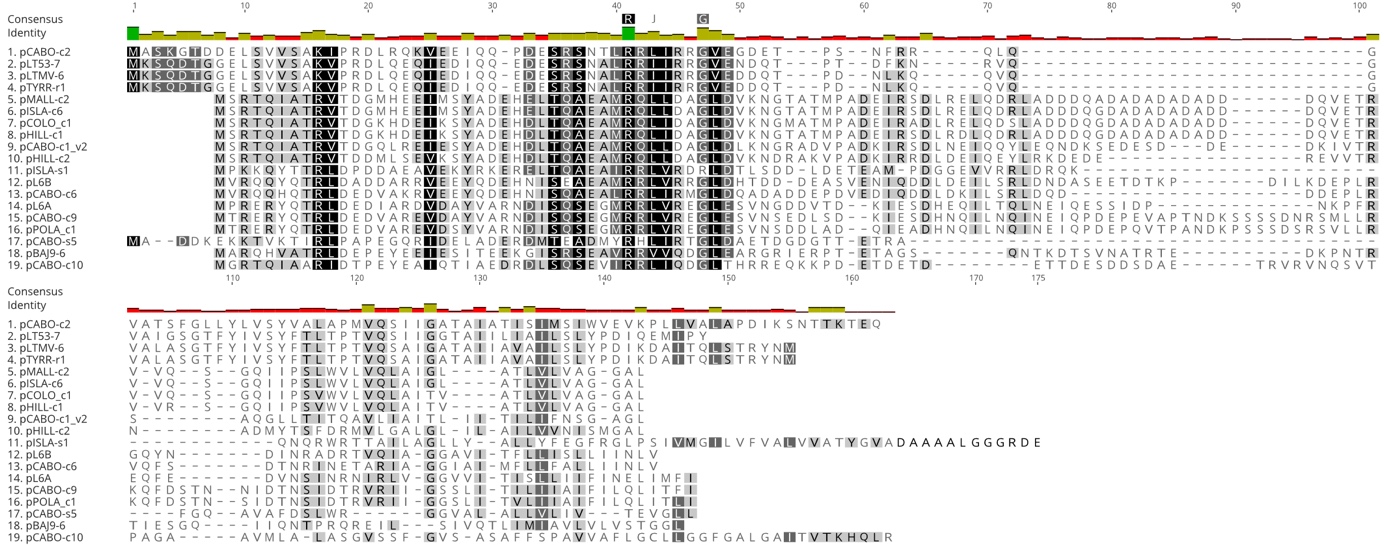

^a^ Inferred R4 protein sequences were aligned according to legend under Figure S2A (above).

**Figure S2, G.** Alignment of R5 proteins^a^
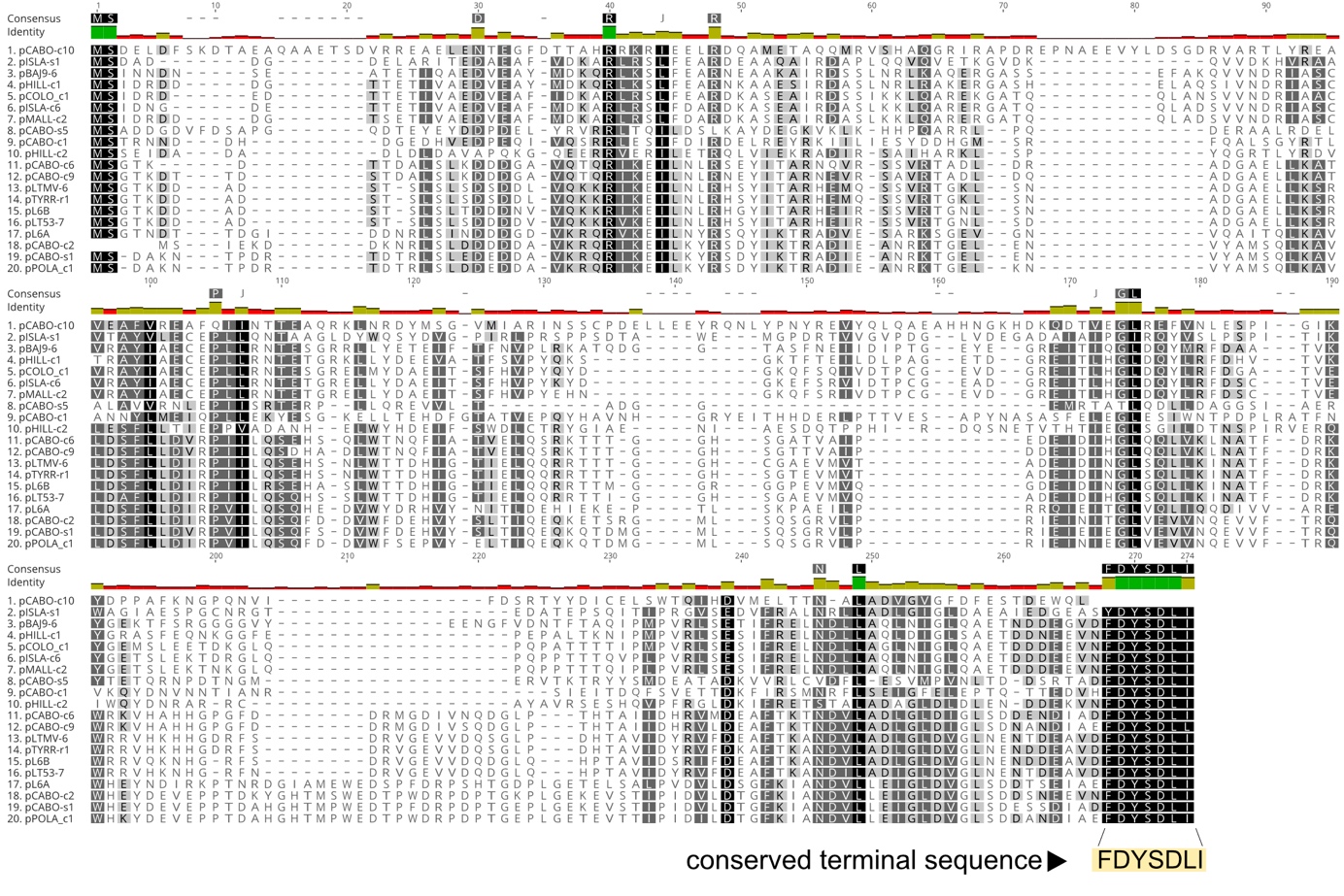

^a^ Inferred R5 protein sequences were aligned according to legend under Figure S2A (above). Note the highly conserved C-terminal sequence [F/Y]DYSDLI.

**Figure S2, H.** Alignment of R6 (ATPase) proteins^a^

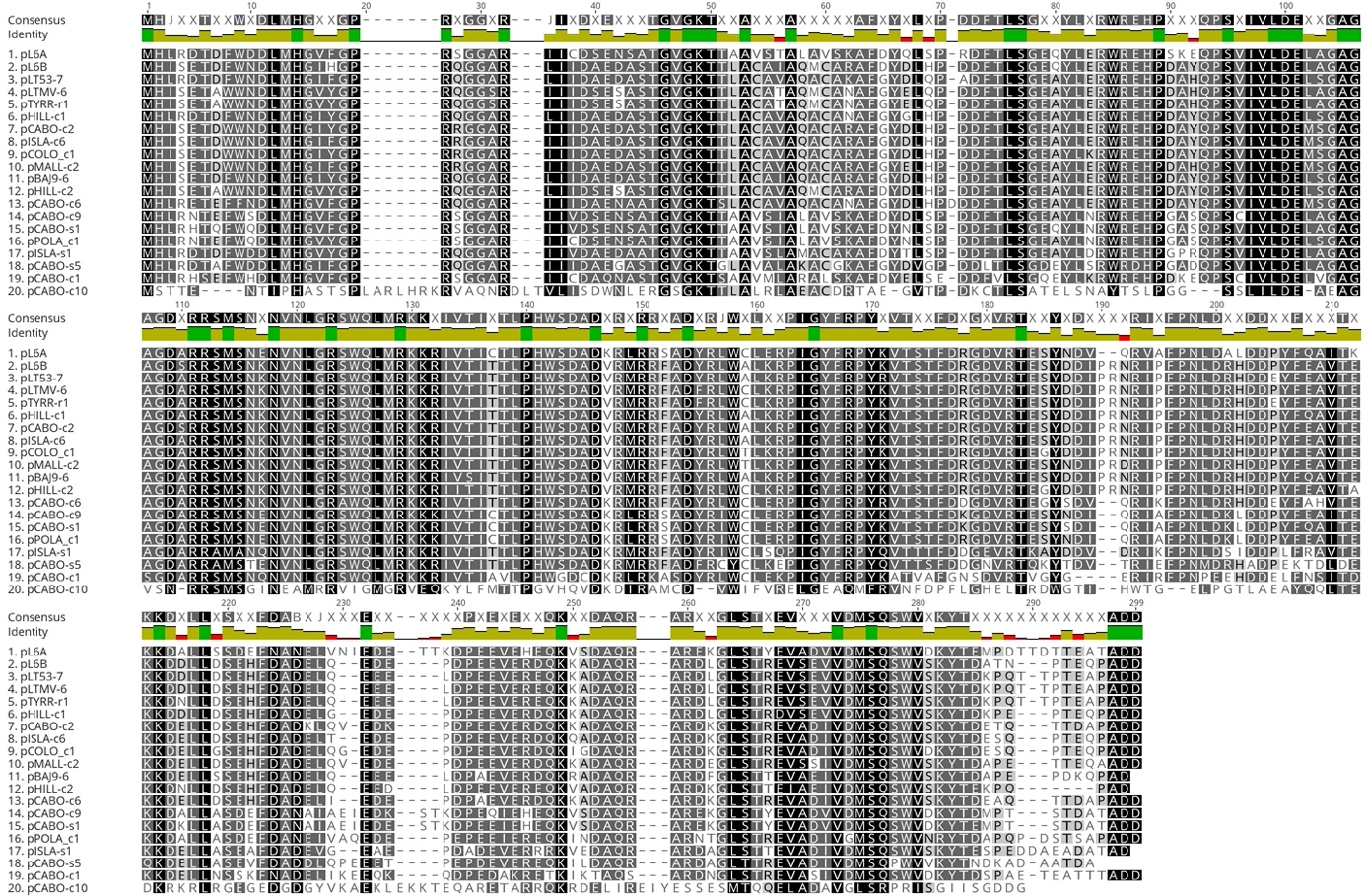

^a^ Inferred R6 protein sequences were aligned according to legend under Figure S2A (above).

**Figure S2, I**. Alignment of R7 proteins^a^

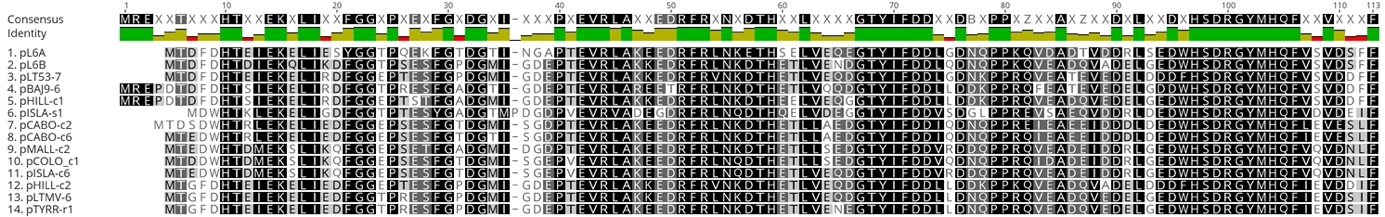

^a^ Inferred R7 protein sequences were aligned according to legend under Figure S2A (above).
