## Supplementary Figures S3 - S7 for "Global distribution and diversity of haloarchaeal pL6-family plasmids"

**Figure S3**. Structural comparison of F2 and F2a proteins

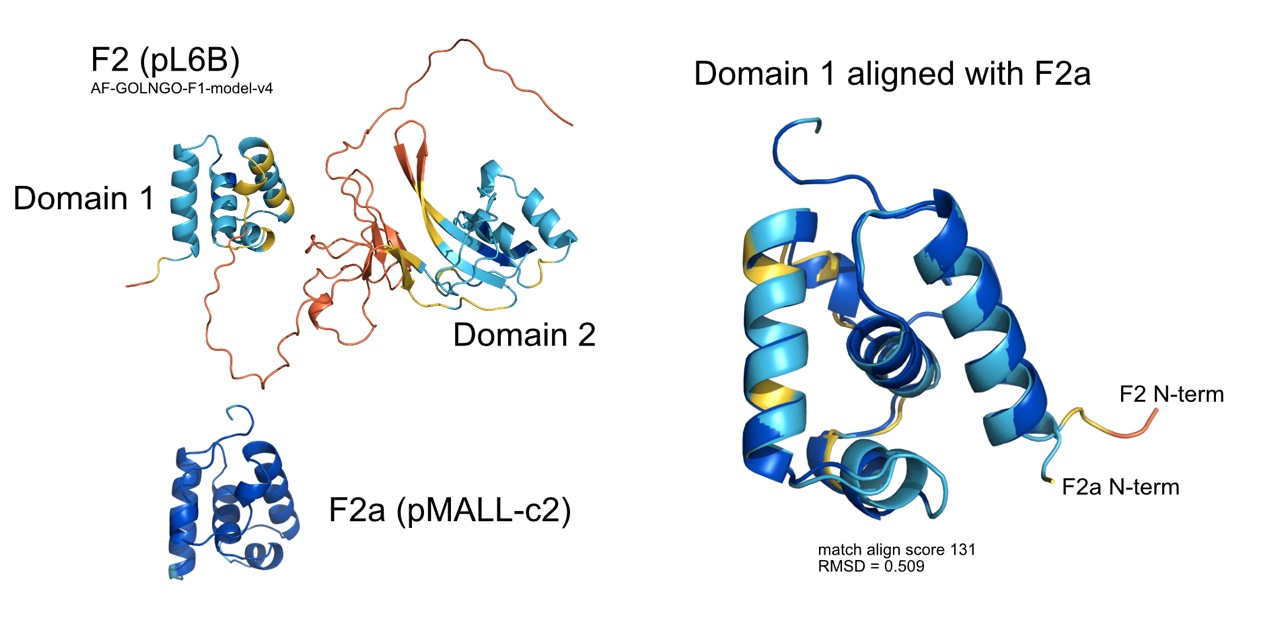

*Upper left*: The alphafold2 (AF2) predicted structure of F2 protein of pL6B is shown (https://alphafold.ebi.ac.uk/entry/G0LNG0) with the standard AF2 coloring denoting prediction confidence (see legend to Figure S27). The two domains are labeled and are separated by a flexible linker. The C-terminus is also flexible.

*Lower left*: AF2 predicted structure of the F2a protein of pMALL-c2.

*Right panel*: Superposition of Domain 1 of F2 and entire F2a protein showing very close structural similarity (match align score 131, RMSD = 0.509)

**Figure S4**. AlphaFold2 prediction of F2b structure (pPOLA-c1)

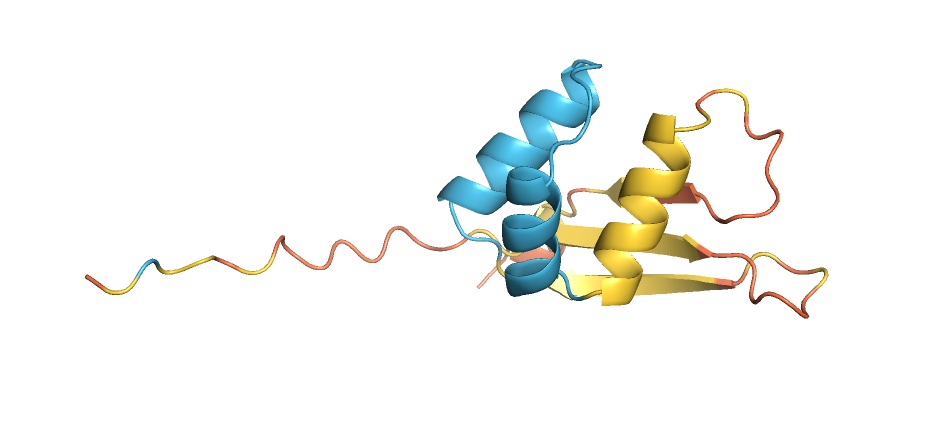

Alphafold 2 prediction via the galaxy server (https://usegalaxy.org.au/). Standard AF2 colouring denoting prediction confidence (see legend to figure S27). N-terminus is at left.

**Figure S5.** AlphaFold 2 structure prediction of halovirus His1_gp16 (AAQ13731)

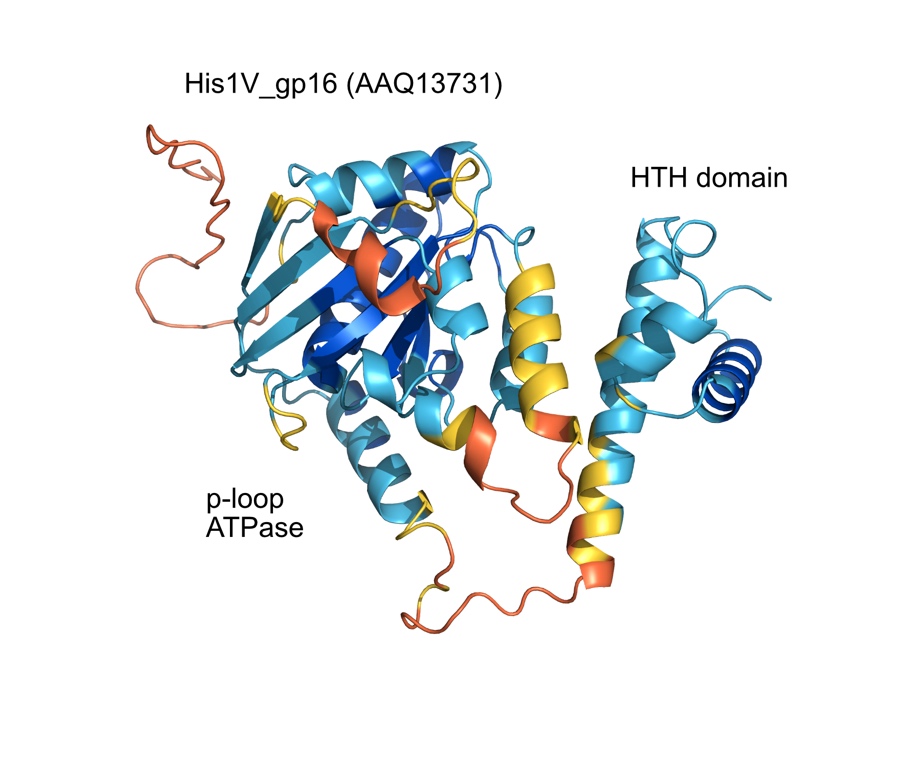

Alphafold 2 predicted structure of halovirus His1 protein His1V_gp16 (AAQ13731). Conserved domains are labeled. Standard AF2 colouring denoting prediction confidence (see legend to figure S27). N-terminus is at left.

**Figure S6.** Comparison of R6 proteins from pL6A and pCABO-c10

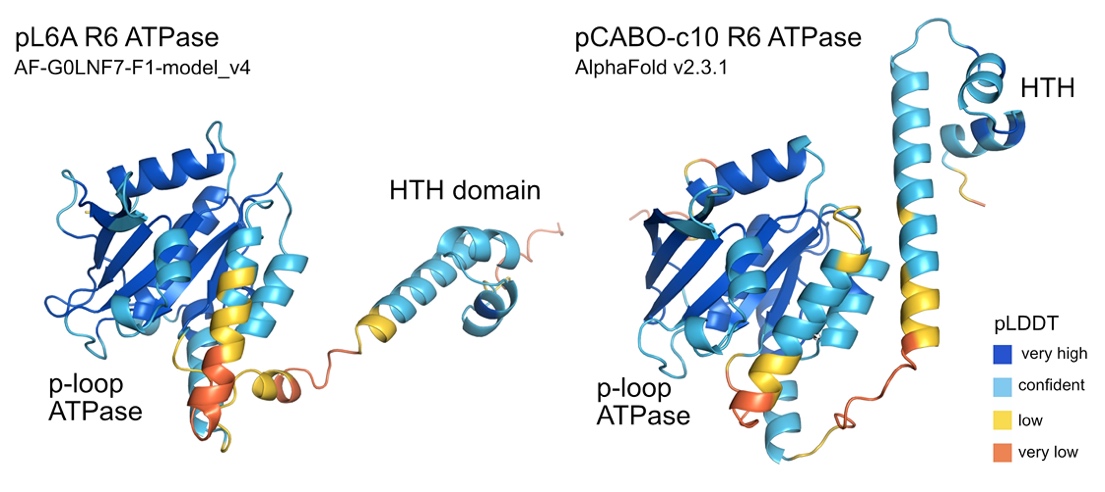

Alphafold 2 predicted structures of R6 proteins of pL6A (left) and pCABO-c10 (right). At lower right is the color code for the standard AF2 colouring denoting prediction confidence.

**Figure S7.** Close structural similarity between formyltransferase encoded by pHILL-c2 and *Mycobacterium tuberculosis* dTDP-4-amino-4,6-dideoxyglucose formyltransferase (P9WKZ3)

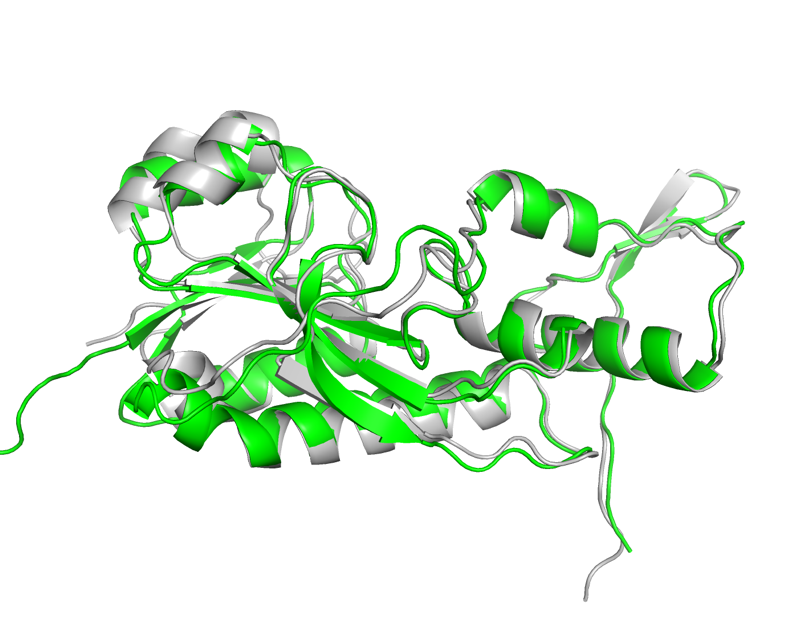

Green chain is the AlphaFold 2 predicted structure (see Methods) of pHILL-c2 formyltransferase. Grey chain is dTDP-4-amino-4,6-dideoxyglucose formyltransferase of *M. tuberculosis* (UniProt accession, P9WKZ3; PDB entry 4PZU) The protein structures were aligned using the align to molecule tool within Pymol v. 2.4.0. The RMSD = 0.794. Substrate binding is in the central cavity (Girardi *et al.*, 2020).
