## Supplementary Figure S8, panels A-C for "Global distribution and diversity of haloarchaeal pL6-family plasmids"

**Supplementary figures S8, panels A-C.**

**Figure S8 A.** Gene neighbours of *T. thiocyanaticus* SAM-dependent methyltransferase WP_125180990.

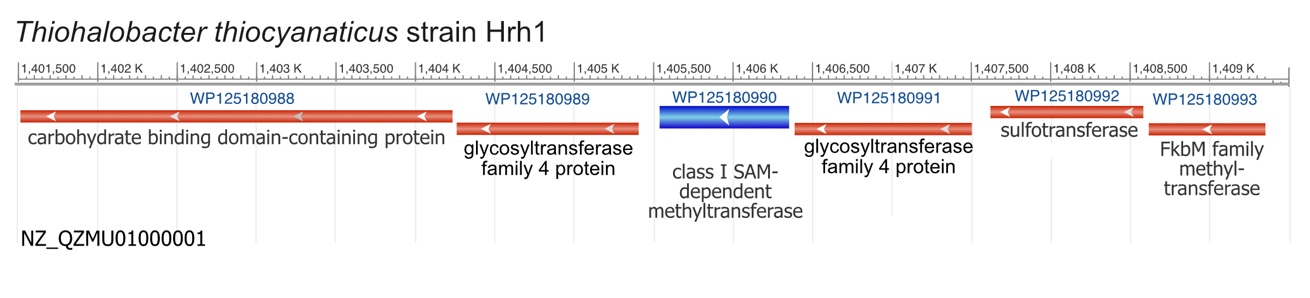

Protein WP_125180990 of *Thiohalobacter thiocyanaticus* strain Hrh1 is a close BLASTp match (40.4% aa identity) to the class I SAM-dependent methyltransferase encoded by pCABO-c9 (Table S4). The gene specifying WP_125180990 (blue bar, centre) is shown along with its gene neighbors on chromosomal contig NZ_QZMU01000001. Note also the FkbM family methyltransferase at the right end, coding for a protein related to the product of the accessory gene of pLT53-7 (see main text). Coordinates are shown in the scale at the top. Coding sequence (CDS) are shown as colored bars (red or blue) and have arrowheads indicating their orientation.

**Figure S8 B.** Gene neighbours of *Har. marina* DT1 FkbM family methyltransferase WP_254272245.1.

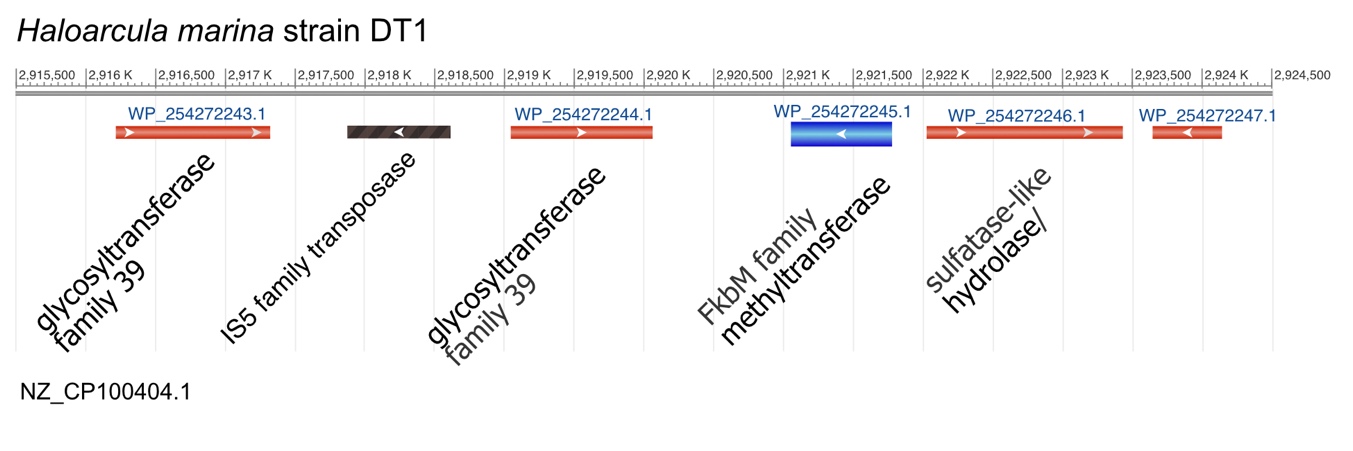

Protein WP_254272245 of *Har. marina* DT1 is a close BLASTp match to pLT53-7 FkbM family methyltransferase (Table S4). The gene encoding WP_254272245 is shown as a blue bar and the surrounding genes are labeled. See legend to figure above for other details.

**Figure S8 C.** Gene neighbours of *Hbt. salinarum* UC4238 gene encoding formyltransferase MDL0133728 (WP_285266011).

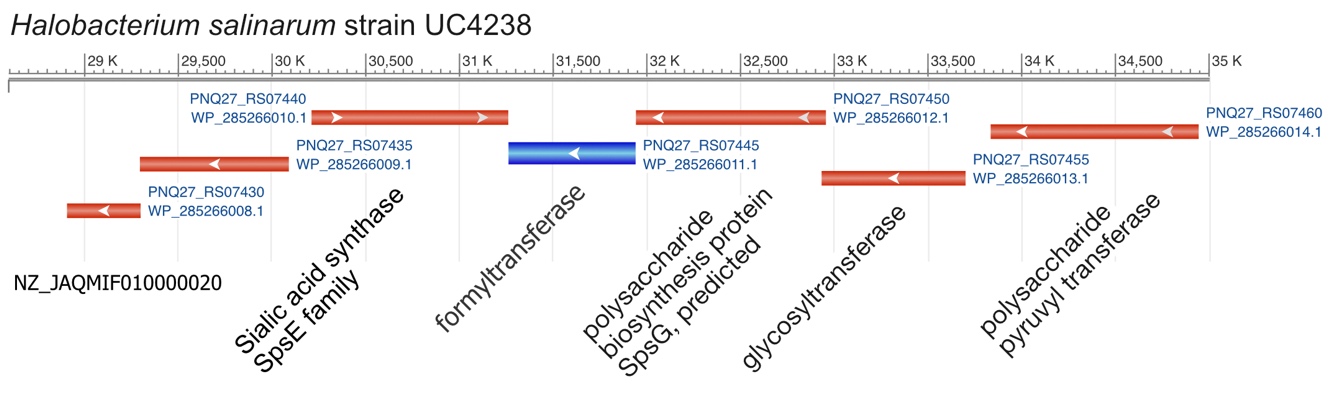

Protein WP_285266011 (MDL0133728) of *Hbt. salinarum* UC4238 is a close BLASTp match to pHILL-c2 formyltransferase. The gene encoding WP_285266011 is depicted by the blue bar in the centre of the gene map (contig ID shown at lower left), along with the surrounding genes (red bars) and the proteins they specify. Contig coordinates shown at the top.
