## Supplementary Figures S9 and S10 for "Global distribution and diversity of haloarchaeal pL6-family plasmids"

**Supplementary figures S9 and S10.** Gene map comparisons.

**Figure S9**. Comparison of two integrative elements containing R6-homologs
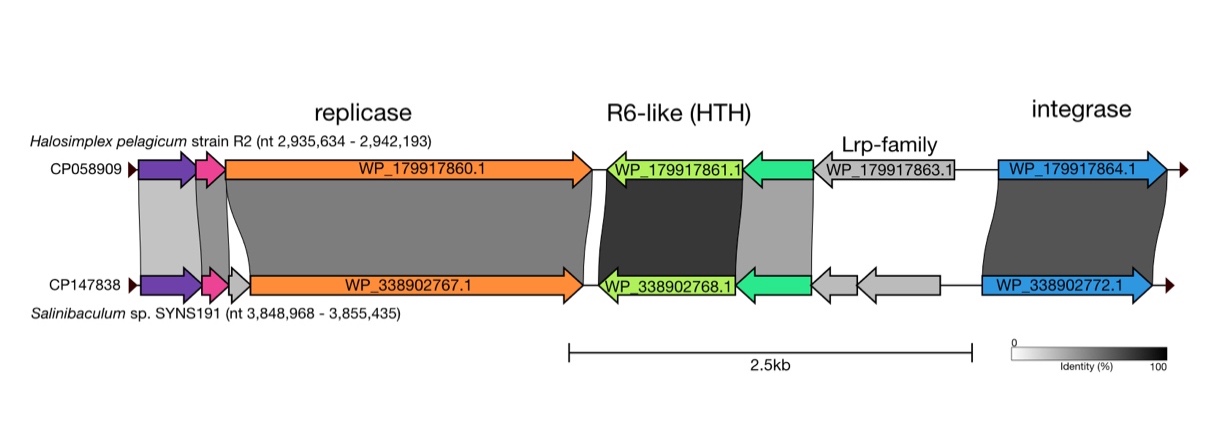

Shaded bands between gene maps represents nucleotide similarity, with values given within or nearby. NCBI accessions are shown at left, and the species names are nearby along with the chromosome coordinates. Small arrowheads at the beginning and end of each gene map represent the direct repeats that flank these elements after integrating at the end of a tRNA-Ala gene. For convenience, several genes are labeled with the accessions for the proteins they specify. The R6-like genes are colored light green (centre). The encoded replicase proteins are 50-53% identical to the viral replicase (ADB79717.1) carried by Haloarcula hispanica pleomorphic virus 1 (HHPV1). In their circular forms, the first CDS (purple) extends leftwards across the direct repeat to begin 3 nt downstream of the end of the integrase gene. See text for other details.

**Figure S10.** Comparison of plasmids pLTMV-6 and pTYRR-r1

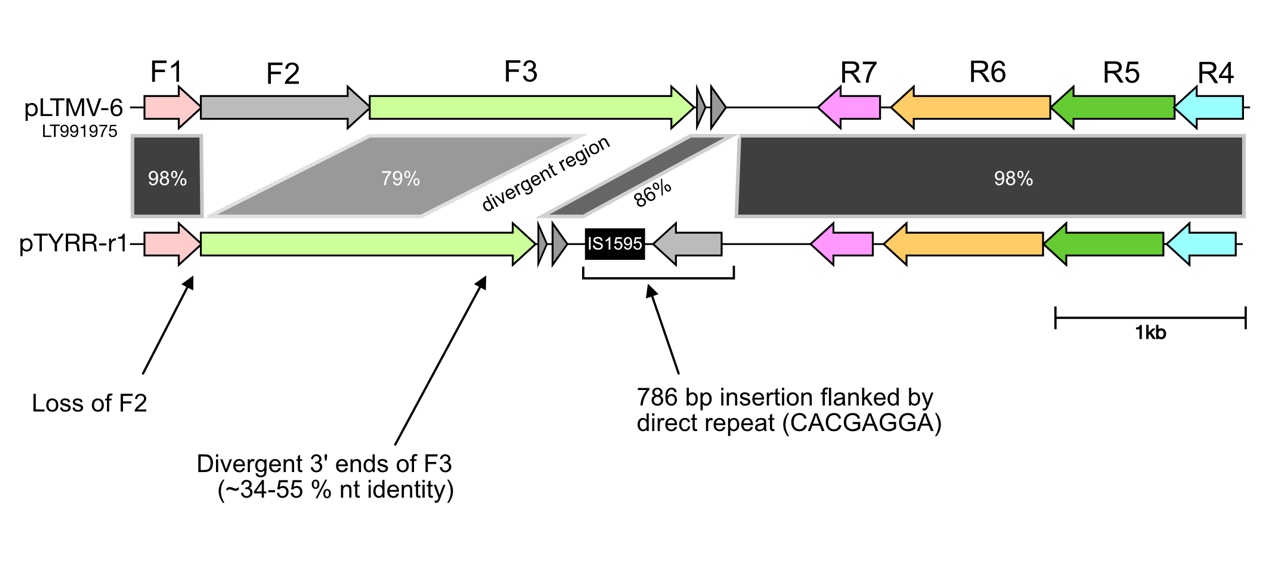

Shaded bands between gene maps represents nucleotide similarity, with values given within or nearby. There is no F2 gene in pTYRR-r1, but the insertion between genes F3 and R7 of pTYRR-r1 means that pLTMV-6 and pTYRR-r1 are of similar size (5,882 and 5,844 bp, respectively).
