## Supplementary Table S1 for "Global distribution and diversity of haloarchaeal pL6-family plasmids"

**Supplementary Table S1. Metagenome, metavirome and RNA-seq read data used in this study**

| Country | Site | Year^a^ | Reads^b^ | Accession^c^ (sample/type) | Reference^d^ |
| --- | --- | --- | --- | --- | --- |
| Argentina | Salina Laguna Colorada Chica | 2019 | 2 × 150 nt | ERR7916263 | (Viver *et al.*, 2023) |
| Australia | Lake Hillier | 2015 | 2 × 250 nt | SRR26978067  (FW-E3) SRR26968577  (FW-D3) | (Sierra *et al.*, 2022) |
| Australia | Lake Tyrrell | 2009 2009 2010 2010 | Average  = 515 nt | SRR5637210 SRR5637211  SRR402042 SRR402044 (virus concentrates) | (Podell *et al.*, 2014) |
| Australia | Lake Tyrrell | 2018 | 2 × 100 nt | SRR24125903-4 (RNA-seq) | (Le Lay *et al.*, 2023) |
| Puerto Rico | Cabo Rojo Saltern | 2014 | 2 × 250 nt | SRR8816317 (sample 1) | (Couto-Rodriguez and Montalvo-Rodriguez, 2019) |
| Puerto Rico | Cabo Rojo Saltern | 2016 | 2 × 150 nt | SRR8816318 (sample 2) SRR8816319  (sample 3) | (Couto-Rodriguez and Montalvo-Rodriguez, 2019) |
| Spain | Isla Cristina saltern | 2020 | 2 × 150 nt | SRR23092357 SRR21894959 | (Garcia-Roldan *et al.*, 2023) |
| Spain | Alicante, Santa Pola saltern, CR30 | 2019 | 2 × 250 nt | SRR13926770 | (Aldeguer-Riquelme *et al.*, 2021) |
| Spain | Mallorca, s'Avall solar saltern | 2018 | 2 × 150 nt | ERR5979340 | (Viver *et al.*, 2023) |
| Spain | Alicante, Santa Pola brines | 2019 | 2 × 100 nt | SRR10674835-40 (RNA-seq) | PRJNA595057 Reg. date: 12-Dec-2019 Centro de Astrobiologia |

^a^ Year of sampling or year of release of sequence data if sampling date is not given.

^b^ Most reads are paired (2 ×). Read lengths given in nucleotides (nt). RNA-seq reads are strand-specific.
^c^Accessions from ENA (https://www.ebi.ac.uk/ena/browser/home) or SRA (https://www.ncbi.nlm.nih.gov/sra).
^d^Either a publication or an NCBI BioProject ID (https://www.ncbi.nlm.nih.gov/bioproject).
