## Supplementary Table S2 for "Global distribution and diversity of haloarchaeal pL6-family plasmids"

**Supplementary Table S2. Mapping of reads to reconstructed pL6-like plasmids.**

| **Plasmid^a^** | **Size (bp)** | **Reads**^b^ | **Reads mapped**^c^ **(threshold %)** | **Coverage**  **(Fold)** | **Total Reads** | **Similarity (%)**^d^ |
| --- | --- | --- | --- | --- | --- | --- |
| pCOLO-c1 | 5,857 | ERR7916263 | 1,744 (0%) | 42 | 72,140,396 | 100 |
| pHILL-c1 | 6,187 | SRR26978067 + SRR26968577 | 4,213 (1%) | 147 | 56,053,528  + 17,295,462 | 99.7 |
| pHILL-c2 | 7,625 | SRR26978067 | 580 (1%) | 18 | 17,295,462 | 99.8 |
| pTYRR-r1 | 5,844 | SRR5637210 | 391 (1%) | 28 | 1,052,913 | 99.3 |
| pCABO-c1 | 5,007 | SRR8816317 | 1,504 (0%) ----------------  2,876 (1%) | 74 -----------  142 | 28,870,606 ------------ 28,870,606 | 100 --------  99.7 |
| pCABO-c2 | 5,889 | SRR8816317 | 1122 (1%) | 47 | 28,870,606 | 99.7 |
| pCABO-c6 | 5,864 | SRR8816317 | 1177 (1%) | 50 | 28,870,606 | 99.7 |
| pCABO-c9 | 6,930 | SRR8816317 | 556 (1%) | 20 | 28,870,606 | 99.7 |
| pCABO-c10 | 5,219 | SRR8816317 | 772 (1%) | 37 | 28,870,606 | 99.7 |
| pCABO-s1 | 6,206 | SRR8816317 | 998 (1%) | 40 | 28,870,606 | 99.7 |
| pCABO-s5 | 5,017 | SRR8816317  + SRR8816318 | 1,053 (1%) | 35 | 28,870,606 25,000,000 | 99.9 |
| pISLA-c6 | 5,881 | SRR21894959 SRR23092357 | 1,327 (1%) | 33 | 145,294,990 + 99,692,172 | 99.7 |
| pISLA-s1 | 5,606 | SRR23092357 | 1,809 (1%) | 47 | 99,692,172 | 99.8 |
| pMALL-c2 | 5,504 | ERR5979340 | 1510 (1%) | 48 | 81,576,380 | 99.8 |
| pPOLA-c1 | 6,075 | SRR13926770 | 1,415 (0%) | 59 | 8,866,062 | 100 |

^a^ Plasmid names with suffix -c were assembled by contig extension; a sequence from an assembled contig was extended by rounds of repeated read mapping (see Methods). The suffix -s refers to plasmids assembled using metaplasmidspades (see Methods). The suffix -r in pTYRR-r1 indicates the initial contig was assembled from RNA-seq reads, and then extended by repeated rounds of read mapping to metagenomic reads.

^b^ NCBI SRA read accessions are given. These were downloaded via the ENA (www.ebi.ac.uk/ena), trimmed and size filtered (see Methods), producing the number of reads shown in the Total Reads column.

^c^ Number of reads mapped to the reference sequence at a maximum level of base differences shown in parentheses. In most cases a base difference tolerance of up to 1% was applied, while in some cases it could be set to zero % mismatches and the coverage still remained high. For pCABO-c1, values are given for both of the applied stringencies. The Geneious mapper (Geneious Prime version 2023.2.1) was set to custom sensitivity, fine tuning = none, minimum mapping quality = 40, only map paired reads which = both map, maximum mismatches per read = 1%, maximum ambiguity = 1.

^d^ Average percentage identity of mapped reads to the reference sequence, as calculated by the Geneious mapper.
