## Supplementary Table S3 for "Global distribution and diversity of haloarchaeal pL6-family plasmids"

**Table S3**. Absent or under-represented tetramers in pL6-plasmids^a^

|  | **Plasmid**^b^ | **Size (bp)** | **%G+C** | **Absent tetramers** | **Under-represented tetramers**^c^ |
| --- | --- | --- | --- | --- | --- |
| Previously published plasmids | pL6A | 6,129 | 51.1 | GGCC | CTAG^3^ |
|  | pL6B | 6,056 | 52.0 | GGCC, CTAG |  |
|  | pBAJ9-6 | 6,213 | 53.0 |  | GGCC^4d^, CTAG^1^ |
|  | pLT53-7 | 7,045 | 50.5 | GGCC | CTAG^2^ |
|  | pLTMV-6 | 5,882 | 53.3 | GGCC, CTAG |  |
| Plasmids reconstructed in the current study | pCABO-c1 | 5,007 | 48.4 | GGCC | CTAG^4^ |
|  | pCABO-c10 | 5,219 | 51.4 | GGCC, CTAG |  |
|  | pCABO-c2 | 5,889 | 53.7 | GGCC, CTAG |  |
|  | pCABO-c6 | 5,864 | 53 | GGCC, CTAG |  |
|  | pCABO-c9 | 6,930 | 49.3 | GGCC | CTAG^1e^ |
|  | pCABO-s1 | 6,206 | 47.8 | GGCC, CTAG |  |
|  | **pCABO-s5** | 5,017 | **58.5** | - |  |
|  | pCOLO-c1 | 5,857 | 53.9 | GGCC, CTAG |  |
|  | pHILL-c1 | 6,187 | 52.4 | GGCC, CTAG |  |
|  | pHILL-c2 | 7,625 | 51.5 | GGCC |  |
|  | pISLA-c6 | 5,881 | 53.6 | GGCC, CTAG |  |
|  | **pISLA-s1** | 5,606 | **64.0** | TTAA |  |
|  | pMALL-c2 | 5,504 | 52.6 | GGCC, CTAG |  |
|  | pPOLA-c1 | 6,075 | 48.1 | GGCC | CTAG^2^ |
|  | pTYRR-r1 | 5,844 | 51.0 | GGCC | CTAG^2^ |

^a^Tetramer searches were performed using the online tool (http://gscompare.ehu.eus/tools/oligo_frequencies/index.php)

^b^Grey shading indicates plasmids previously reported in (Dyall-Smith and Pfeiffer, 2018). Bold type denotes the high GC plasmids.

^c^Under-represented tetramers have a superscript number for the tetramer count.

^d^Four GGCC motifs occur within two closely spaced repeats found in the accessory gene region of this plasmid.

^e^The single CTAG site occurs in a foreign gene (SAM-dependent methyltransferase)
